# An Atypical E3 Ligase Module in UBR4 Mediates Destabilization of N-degron Substrates

**DOI:** 10.1101/2023.05.08.539884

**Authors:** Lucy Barnsby-Greer, Peter D. Mabbitt, Marc-Andre Dery, Daniel R. Squair, Nicola T. Wood, Sven Lange, Satpal Virdee

## Abstract

UBR4 is an E3 ligase (E3) of the N-degron pathway and is involved in neurodevelopment, age-associated muscular atrophy and cancer progression. The location and mechanistic classification of the E3 module within the 600 kDa protein UBR4 remains unknown. Herein, we identify and characterize, at a biochemical and structural level, a distinct E3 module within human UBR4 consisting of a novel “hemiRING” zinc finger, a helical-rich UBR Zinc-finger Interacting (UZI) subdomain, and a predicted backside interacting N-terminal helix. A structure of an E2 conjugating enzyme (E2)-E3 complex provides atomic level insight into the exquisite specificity of the hemiRING towards the E2s UBE2A/B. The UZI subdomain can be considered a component of the E3 module as it has a modest activating effect on the ubiquitin loaded E2 (E2∼Ub), which is complemented by the intrinsically high lysine reactivity of UBE2A. These findings reveal the mechanistic underpinnings of a neuronal N-degron E3 ligase, its specific recruitment of UBE2A, and highlight the underappreciated architectural diversity of cross-brace domains associated with ubiquitin E3 activity.

## Main

The N-degron and C-degron pathways are fundamental aspects of the ubiquitin (Ub) system ^1^. These pathways regulate protein stability based on the identity of either the N- or C-terminal amino acid of a protein substrate where destabilization is mediated by proteasomal or autophagic clearance. The regulated degradation of substrates by the N-degron pathway affects multiple cellular processes including elimination of misfolded or mislocalized proteins, maintenance of protein complex stoichiometry, DNA repair, apoptosis, metabolite sensing, and neurodevelopment ^1^. N-degrons are recognized by N-recognins, which are typically ubiquitin E3 ligases (E3s). Type 1 N-degrons are strongly destabilizing and are recognised by a dedicated ∼70 residue zinc finger domain known as the UBR-box ^2, 3^.

In mammals, there are 7 UBR family members (UBR1-7). The E3 module responsible for ubiquitination activity in UBR1-3 is the RING domain ^1, 2, 4^. The RING domain is a small (∼10 kDa) fold that coordinates two Zn^2+^ ions with a cross-brace configuration ^5^. The U-box domain achieves a similar fold through hydrogen-bonding networks ^6^. RING/U-box E3s catalyse Ub transfer from a thioester-linked E2 conjugating enzyme (E2∼Ub) to substrates. They use an allosteric “adapter-like” mechanism involving protein-protein interactions rather than a defined active site ^7–11^. This catalyzes substrate ubiquitination by stabilizing a closed E2∼Ub conformation which has enhanced lysine reactivity ^9–13^. However, RING domains can also have a more benign function as a docking site for E2∼Ub and this is exemplified by RING-in-between-RING (RBR) and RING-Cys-Relay (RCR) E3s ^14, 15^. Both contain an ancillary domain harboring an essential active site cysteine that undergoes a transthiolation reaction with E2∼Ub. Similarly, Homologous to E6AP Carboxy-Terminus (HECT) E3s also have an active site cysteine in their HECT domain and UBR5 belongs to this class ^16^. There is further mechanistic divergence within the UBR family as UBR6 (more commonly known as Fbxo11) is a substrate receptor of a multi-subunit Cullin RING E3 complex^17^, and UBR7 contains a plant homeodomain (PHD) that possesses E3 activity ^18, 19^.

UBR4 is an essential protein ubiquitously expressed in all tissues but is highly enriched in the central nervous system (CNS) and has several prominent functions in the mammalian brain including neurogenesis, neuronal migration, neuronal survival and signalling ^20–24^. UBR4 has also been implicated in anoikis, viral transformation and the endosome-lysosome system ^25, 26^. Disease associations include neurological disorders and myofiber atrophy ^27–32^, and UBR4 loss increases cancer cell susceptibility to apoptosis ^33^. The latter might be due to proteotoxicity from imbalances in protein complex stoichiometry arising from aneuploidy - a hallmark of many cancers. Mechanistically, little is known about UBR4 and none of its domains have been structurally characterized. Furthermore, UBR4 does not contain a predictable E3 module therefore the source of its E3 activity remains a mystery. This highlights a notable gap in our molecular understanding of this protein and the N-degron pathway in general.

## Results

### A C-terminal UBR4 E3 module is catalytically functional with UBE2A/B

UBR4 is one of the largest known single subunit proteins consisting of over 5,000 residues (**Fig. 1a**) ^25^. To establish the mechanistic subtype and location of the E3 module we initially tested which E2s support UBR4 E3 activity. Although UBR4 has been shown to bind the E2 UBE2A, this interaction has not been functionally tested ^34^. The E2 activity profile of UBR4 could also provide insight into whether it utilizes an adapter-like or transthiolating mechanism ^14^. UBR4 is a large multidomain protein so to ensure comprehensive assessment of E3 activity we immunoprecipitated full-length HA-tagged UBR4 we stably expressed in HEK293 cells. Autoubiquitination activity of HA-UBR4 was then measured with a recombinant panel of 29 ubiquitin E2s (**Extended Data Figure 1**). HA-UBR4 underwent robust autoubiquitination activity but only when partnered with UBE2A or its paralogue UBE2B (**Fig. 1b**). UBE2A (also known as RAD6A) has been reported to cooperate with adapter-like E3s and further suggestive of UBR4 belonging to this mechanistic subtype, autoubiquitination was absent with UBE2L3 as this E2 is understood to only support transthiolation E3 activity ^14^.

**Figure 1.**
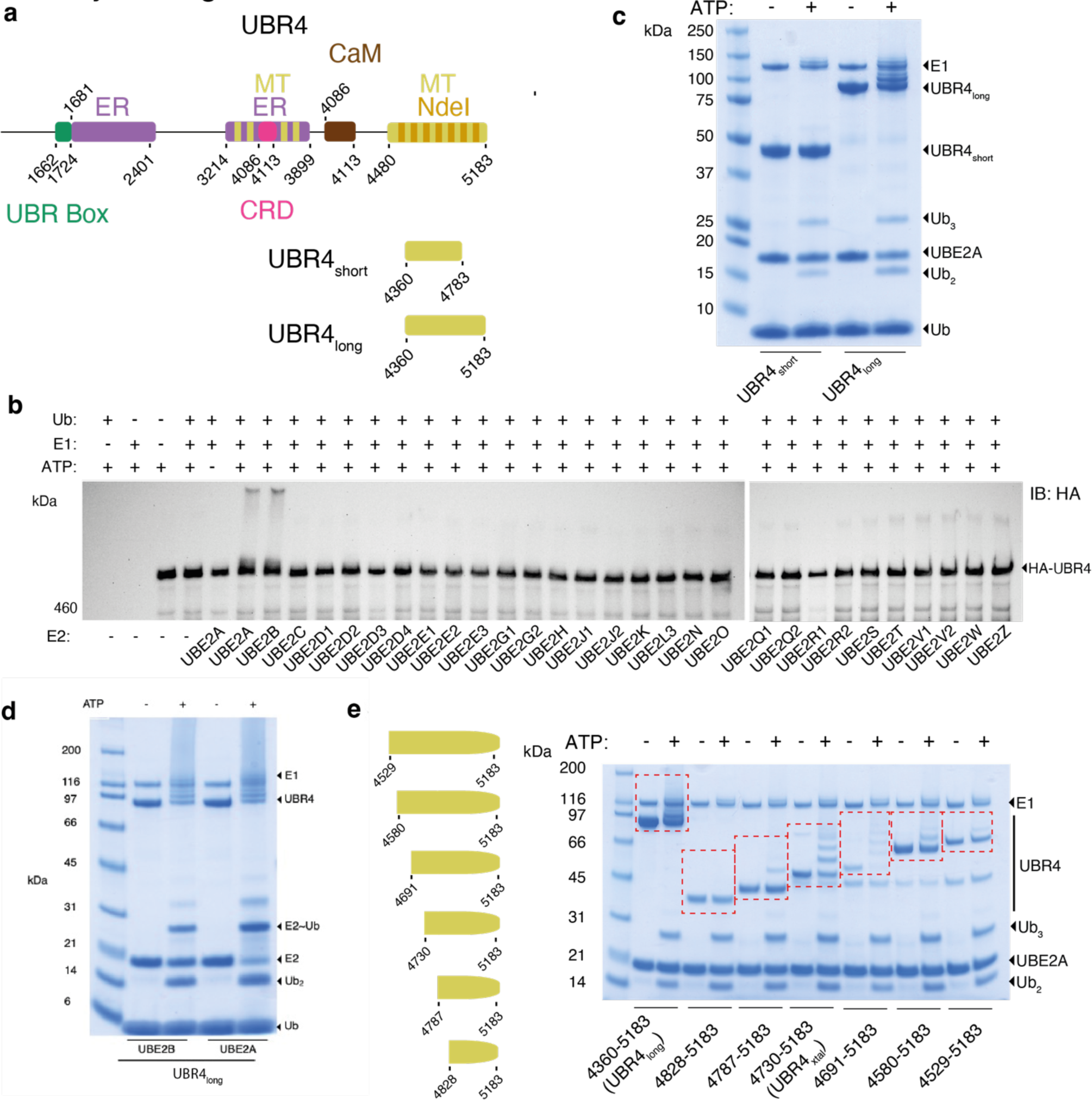
Domain Architecture of UBR4 and assessment of UBR4 E3 ligase activity. **a)** Domain architecture of UBR4. Abbreviations defined as follows: UBR-box domain, Endoplasmic Reticulum-associated region (ER), Microtubule binding region (MT), Cysteine-rich Domain (CRD), Calmodulin-binding Domain (CaM), Ndel1-binding region (Ndel1). Regions in alternating-colored bars correspond to those that have more than one experimentally observed interaction ^21, 25^. Two recombinant scout constructs were used to identify the E3 module, UBR4_short_ and UBR4_long_. **b)** Twenty-nine E2s were screened for their ability to support autoubiquitination activity with full-length UBR4. A stable HEK293 cell line expressing full-length HA-UBR4 was immunoprecipitated using anti-HA sepharose resin. Washed resin was combined with E1 (500 nM), ubiquitin (5 µM), ATP (10 mM) and the specified E2 (1.5 - 10 µM). Reactions were incubated at 37 °C for one hour, stopped by the addition of reducing LDS loading buffer and visualized by anti-HA immunoblot. **c)** When partnered with UBE2A, only UBR4_long_ underwent autoubiquitination whereas no activity was observed with UBR4_short_. Reactions were resolved by SDS-PAGE and visualized by Coomassie staining. **d)** UBR4_long_ demonstrated comparable autoubiquitination activity with UBE2A and UBE2B. **e)** Progressive N-terminal truncations of UBR4_long_ were tested for autoubiquitination activity with UBE2A. The UBR4 construct and its autoubiquitination products are highlighted with red hashed boxes.

E3 modules often exist at the C-terminus of their polypeptide sequences so we designed two recombinant scout fragments amenable to expression in *Escherichia coli*. Firstly, a ∼93 kDa construct consisting of the C-terminal 823 residues (UBR4_long_; residues 4360-5183) and a 47 kDa C-terminal truncated version (UBR4_short_; residues 4360-4783), both of which are within the second microtubule binding region of UBR4 (**Fig. 1a**). UBR4_long_ underwent robust autoubiquitination when partnered with UBE2A but lack of activity with UBR4_short_ revealed the functional importance of the C-terminal 400 residues (**Fig. 1c**). UBR4_long_ was also similarly active with UBE2B (**Figure 1d**). Progressive truncation of the N-terminus of UBR4_long_ identified UBR4_4730-5183_ as the minimal construct that retained autoubiquitination activity (**Fig. 1e**). A caveat with autoubiquitination assays is they do not discern whether loss of autoubiquitination is due to catalytic impairment or loss of autoubiquitination sites. Nevertheless, we focused on the UBR4_4730-5183_ construct for further characterization.

### Adapter-like UBR4 E3 activity exploits high UBE2A lysine reactivity

To further corroborate an adapter-like mechanism for UBR4_4730-5183_, we tested activity with two UBE2A mutants. Residue Asn80 is part of a highly conserved His-Pro-Asn motif where the asparagine residue has an essential role in thioester activation and/or transition state stabilization but is typically dispensable for E3s that undergo transthiolation by engaging an open E2∼Ub conformation^9, 14, 35–39^. On the other hand, residue Ser120 (aspartate 117 in UBE2D1-4) can be considered a requirement for E2 transfer to lysine, which is uniquely exploited by adapter-like E3s ^40, 41^. The requirement for these residues was assessed by quantitative gel-based autoubiquitination assays utilizing fluorescent Cy3b-labelled Ub. To ensure potential perturbation to E1-E2 activity was decoupled, assays were carried out under single turnover E2∼Ub discharge conditions ^15^. We found that neither the Asn80Ser nor the Ser120Ala mutant could support UBR4 autoubiquitination activity (**Fig. 2a, b**). Taken together, these data are consistent with the E3 module residing within UBR4 residues 4783-5183 operating via an adapter-like mechanism.

**Figure 2.**
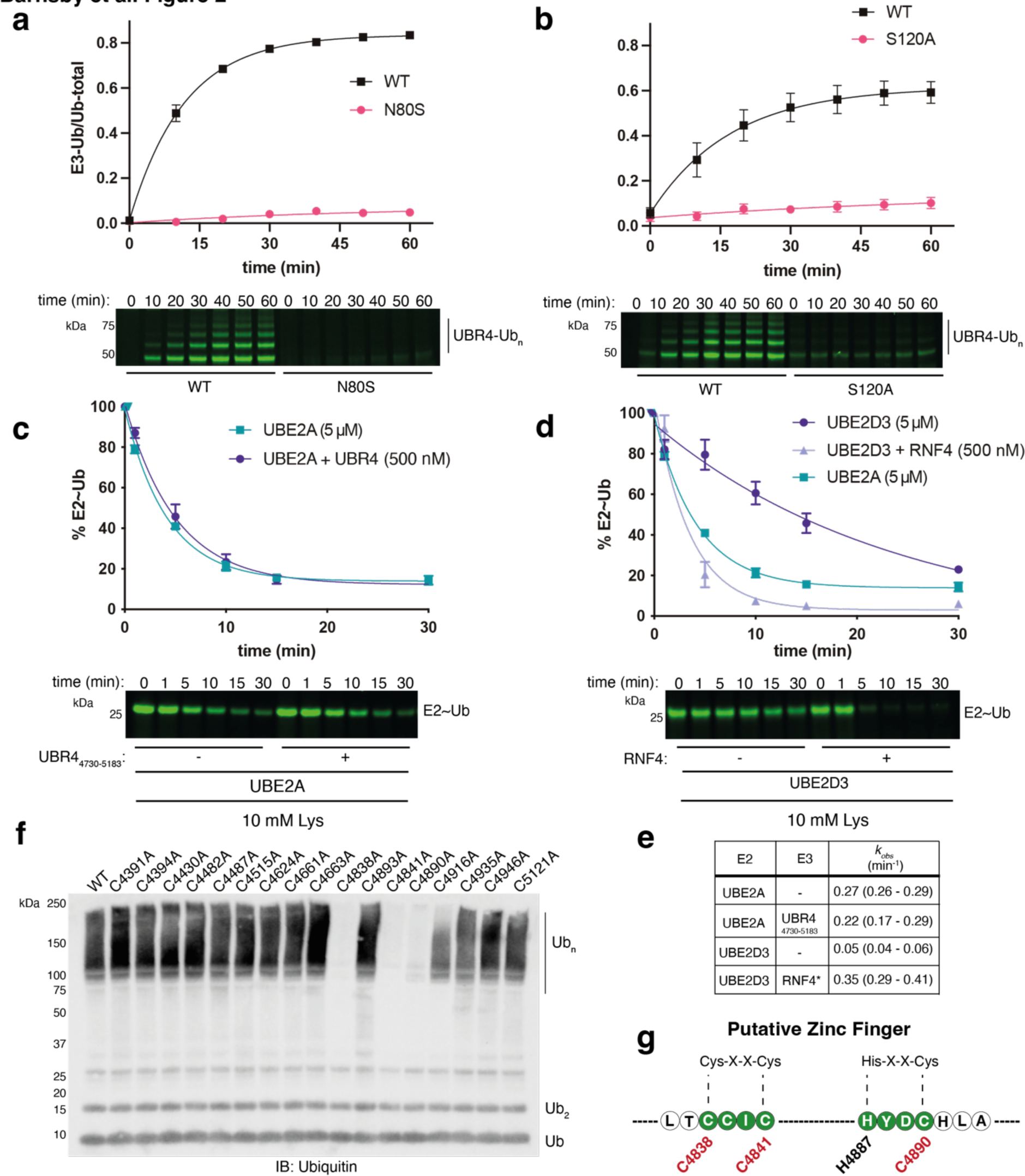
Biochemical activity profiling reveals the presence of an adapter-like E3 module based on a putative zinc finger near the UBR4 C-terminus. **a)** Quantitative UBR4 autoubiquitination assay comparing UBE2A WT with UBE2A Asn80Ser under single turnover E2∼Ub discharge conditions (*n* = 3 and standard error is displayed). **b)** Autoubiquitination assay comparing UBE2A WT with UBE2A Ser120Ala (*n* = 3 and standard error is displayed). **c)** Single turnover discharge of Ub from UBE2A (5 μM) to lysine in the presence and absence of UBR4_4730-5183_ (500 nM) (*n* = 3 and bars represent the standard error). Activation was also not observed at a higher (5 μM) UBR4 concentration (Extended Data **Fig. 2**). **d)** Single turnover discharge of Ub from UBE2D3 to lysine in the absence or presence of a subsaturating concentration of an RNF4 variant (500 nM) engineered to be constitutively active (*n* = 3 and bars represent standard error) ^42^. For reference, the UBE2A curve for E3-independent lysine discharge from panel c is also presented. **e)** Observed rates determined from the experiments presented in c and d using the half-life equation (*k_obs_* = ln2/t_1/2_). Errors correspond to the 95% confidence intervals obtained from fitting the data to a single exponential function. **f)** The seventeen cysteines within UBR4_long_ were systematically mutated to alanine and autoubiquitination activity was assessed by anti-Ub immunoblot. **g)** Cysteine residues required for robust autoubiquitination activity can be ascribed to a putative C2HC zinc finger that binds a single zinc ion.

Associated with thioester activation is the formation of a closed E2∼Ub conformation that is allosterically stabilized by adapter-like E3s ^9–11^. To test for this characteristic, we used free lysine as a model substrate ^14^. However, under our assay conditions UBR4_4730-5183_ did not discernibly stimulate Ub discharge from UBE2A/B, suggesting it does not stabilize the closed E2∼Ub conformation, or at least not to the same extent as prototypical RING E3s (**Fig. 2c** **and Extended Data Fig. 2**) ^9–11^. UBE2A reactivity when partnered with a cognate E3 has not been biochemically investigated in detail so we asked if it possesses intrinsically high lysine aminolysis activity, because this would reconcile the lack of E3-dependent lysine discharge. We determined the observed rate of UBE2A discharge to lysine and under our conditions we found it was at least 6-fold higher than for the prototypical E2 UBE2D3 **(Fig. 2d, e)**. Strikingly, the observed rate of UBE2A discharge was comparable to Ub discharge from UBE2D3 when catalyzed by a constitutively active variant of the prototypical RING E3 RNF4 **(Fig. 2d, e)** ^42^. Thus, lack of, or tempered thioester activation might be characteristic of E3s that are cognate for UBE2A/B because this can be compensated for by their intrinsically high aminolysis activity. Such a model would also account for the ability of UBE2A/B to mediate highly efficient E3-independent proximity-induced protein degradation ^43^.

### UBR4 E3 ligase activity is dependent on a putative zinc finger motif

Zinc finger domains are often associated with E3 activity, and their structural integrity is dependent on cysteine/histidine residues that coordinate zinc ions ^5, 15, 44^. To see if a cryptic zinc finger existed within UBR4 we mutated the 17 cysteines in UBR4_long_ to alanine and assessed the effect on autoubiquitination (**Fig. 2f**). We found mutation of three cysteine residues (C4838A, C4841A, C4890A) abolished activity. Whilst this region remains unannotated, the position of C4838 and C4841 correspond to a Cys-X-X-Cys motif characteristic of a zinc finger ^5^. A single zinc ion is usually coordinated by four residues and H4887 at the -3 position relative to C4890 (His-X-X-Cys) could complete a potential ligand network (**Fig. 2g**). This raised the possibility that these are required for the structural integrity of a cryptic C2HC zinc finger that functions as an E2 docking site. Consistent with this putative structural motif being central to E3 activity, full length proteins immunoprecipitated from HEK293 cells containing H4877A or C4890A mutations were devoid of autoubiquitination activity **(Extended Data Fig. 3**).

### The hemiRING-UZI is a novel E3 module within UBR4

To gain further insights into the unidentified E3 module we solved the structure of UBR4_4730-5183_. We obtained crystals of UBR4_4730-5183_ (referred to hereon as UBR4_xtal_) after sparse matrix screening and condition optimization. Diffraction data were collected and a 1.8 Å structure was solved using the anomalous signal from the zinc ion present in the structure (**Fig. 3a, b and Extended Data Fig. 4, 5**). Residues 4730-4830 of UBR4_xtal_ were not resolved so a model could only be built for residues 4831-5183 (**Fig. 3b**). Of note, a designed construct approximating this region (UBR4_4828-5183_) was inactive in our autoubiquitination assay (**Fig. 1e**). Our crystal structure is made up of two apparent subdomains that constitute a larger fold. The N-terminal subdomain comprises residues 4835-4948 and is followed by the second subdomain composed of 11 alpha helices that runs to the native UBR4 C-terminus (**Fig. 3c**). No established folds with clear homology to this region were identified with the DALI comparison server but the most similar was rhamnosidase B (DALI Z-score 6.5) ^45^. As such we consider this subdomain a novel fold and refer to it as the UBR Zinc-finger Interacting (UZI) region. Interestingly, two mutations found in episodic ataxia patients, Ala5042Val and Arg5091His, reside within this region **(Extended Data Fig. 6a, c)** ^46, 47^. Ala5042 is located within helix α5 and packs against helices α3 and α6 whereas Arg5091 is in helix α7 and forms a strong salt bridge with Glu4971 in the α1-2 loop (**Fig. 3c**). We found that the Ala5042Val mutant exhibited impaired autoubiquitination activity whereas a very modest impairment was observed with Arg5091His **(Extended Data Fig. 6b, d)**.

**Figure 3.**
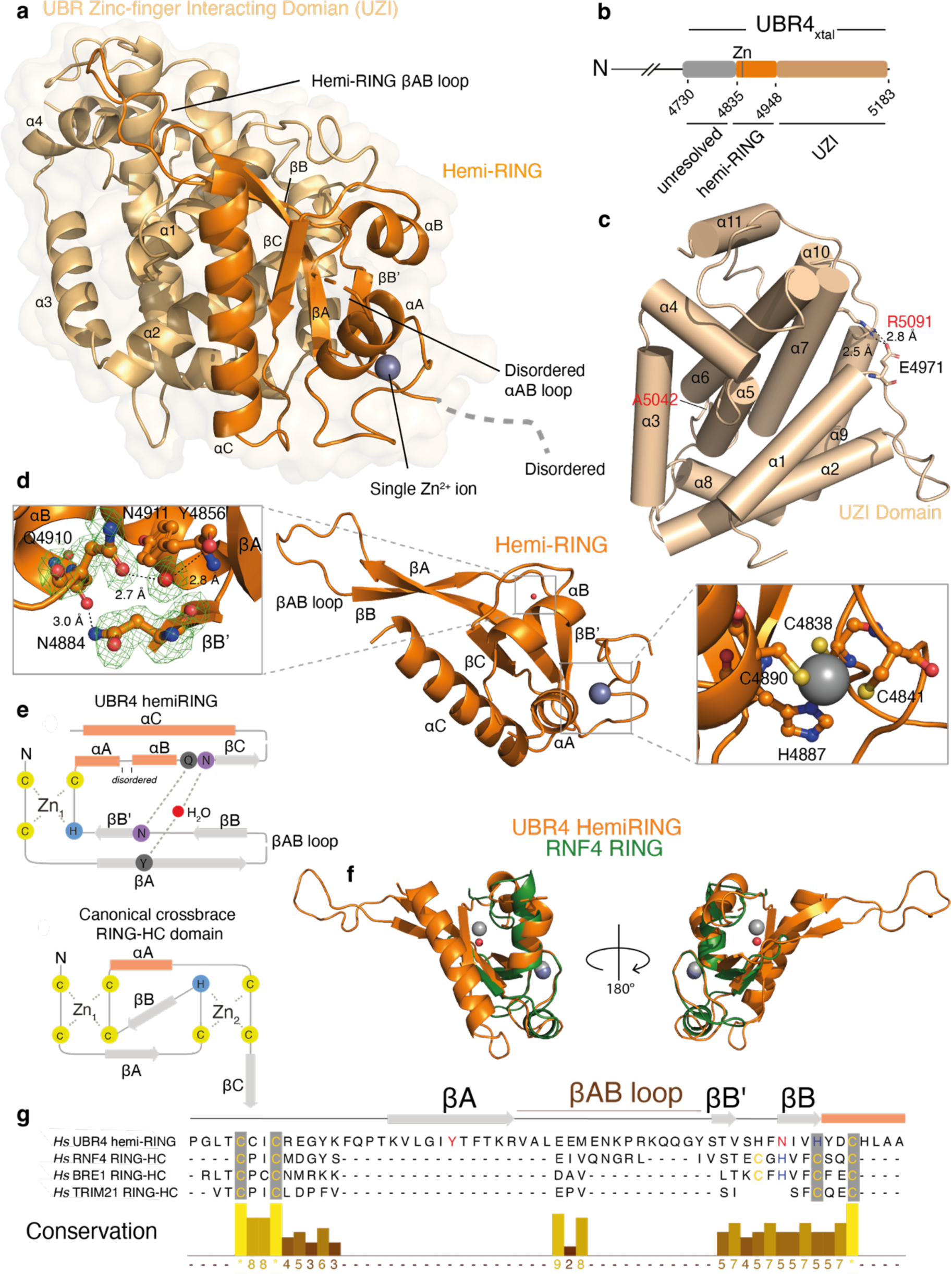
Crystal structure of core zinc finger UBR4 E3 module. **a)** Surface and cartoon representation of the hemiRING-UZI domain structure refined to 1.8 Å. The hemiRING subdomain is colored in orange whereas the UZI domain is colored in wheat. N-terminal residues 4730-4832 and residues 4897-4901, that connect helices αA and αB, were found to be unresolved in the crystal structure. **b**) Domain architecture of hemiRING-UZI subdomain. Color as for **a** but a region in the crystallization construct (UBR4_xtal_) that was not present in the crystal structure is highlighted in grey. **c**) The UZI domain is a novel fold composed of a bundle of 11 α-helices. Residues mutated in episodic ataxia patients are depicted in ball and stick. **d**) Close up of the hemiRING structure. Inset (left) depicts the water-mediated and conventional hydrogen bonding network that apparently substitutes for coordination of a second zinc ion. Green mesh corresponds to a *F_obs_-F_calc_* difference map for Asn884, Gln4910 and water molecule 199 calculated with Phenix and contoured at 3.0 α . Inset (right) depicts the C2HC coordination network for the single zinc ion present in the structure. **e**) Schematic representations of a canonical RING-HC domain and the UBR4 hemiRING. Orange rectangles represent α-helices and grey arrows represent ý-strands. Residues that hydrogen bond through their backbone are in grey circles whereas those that interact through their side chain are in purple circles. **f**) Structural superposition of the UBR4 hemi-RING (orange) onto the RNF4 RING domain (green; PDB 4PPE). The hemiRING Zn^2+^ ion is a dark grey sphere, and a water molecule is a red sphere. The two zinc ions in RNF4 are light grey spheres. **g)** Sequence alignment of the *Homo sapiens* (*Hs*) UBR4 hemiRING sub-module with cross-brace RING domains from *Hs* RNF4, BRE1 and TRIM21. Cysteine and histidine residues that coordinate a zinc ion are highlighted in yellow or blue, respectively. The UBR4 residues that stabilize the cross-brace fold through hydrogen bonding are highlighted in red.

Our experimental structure confirms the structural role of C4838, C4841, C4890 and H4887 as ligands that coordinate a single zinc ion within the zinc finger and these residues are conserved in UBR4 orthologues (**Fig. 3d** **and Extended Data Fig. 5b, 6h)**. Although not evident from primary sequence, the zinc finger has partial structural homology with canonical RING domains (DALI Z-score range 2.0-5.8) **(Fig. 3e, f)**. Although the protein fold immediately proximal to zinc ion 1 (referred to hereon as the proximal Zn site), which engages the E2 enzyme in canonical RING domains is conserved together with the tailing helix and core beta sheet (**Fig. 3e, f**) ^48^, residues essential for coordinating the second zinc ion (referred to hereon as the distal Zn site) are conspicuously absent. Substituting for zinc ion 2 are four residues that form hydrogen bonds and are likely required for stabilizing the cross-brace architecture of the zinc finger subdomain (**Fig. 3d, e**). Tyr4856 forms a water-mediated hydrogen bond via its backbone carbonyl to the side chain of Asn4911, and these are located within strand βA and the end of helix αB, respectively. Following the His-X-X-Cys motif that completes coordination of the single zinc ion the side chain of Asn4884, located within strand βB’, hydrogen bonds with the backbone carbonyl of Gln4910 (**Fig. 3d, e**). Except for Tyr4856, whose side chain is solvent exposed and hydrogen bonds via its backbone carbonyl, these residues are also conserved across orthologues (**Extended Data Fig. 6h**). The hybrid U-Box/RING nature of the UBR4 zinc finger is reminiscent of the SP-RING domain found in SUMO E3 ligases of the Siz/PIAS family ^49^. However, the SP-RING maintains the distal Zn site rather than the proximal site and lacks the extended βAB sheet. Thus, to our knowledge the UBR4 zinc finger represents a novel RING-related fold and we refer to it as the hemiRING.

Interestingly, an extended beta sheet composed of the ýA and ýB strands packs against a helix at the end of the hemiRING and α2 of the UZI domain. The ýA and ýB strands are connected via a lengthy loop, which makes further contacts with helices α1 and α4 of the UZI subdomain (**Fig. 3e, f, g** **and Extended Data Fig. 5**). Another episodic ataxia mutation is Tyr4877Cys and is located within the hemiRING βB’ strand, which makes hydrophobic contacts with the UZI **(Extended Data Fig. 6e**). Despite being highly conserved, its introduction had no discernible effect on autoubiquitination (**Extended Data Fig. 6f, h)**. The RING domain in yeast UBR1 has an even longer loop that is sandwiched by a helical-rich fold known as the cap helical domain (CHD) ^4^. We extended the fold comparison of the UZI subdomain beyond experimentally determined structures using the DALI AF-DB comparison tool ^50^. Strikingly, the top matches were the N-degron E3 ligases from *Homo sapiens* that contain a RING domain (UBR1, UBR2 and UBR3; Z-score 6.6-8.0; RMSD 3.8-4.2 Å). The 11 helices comprising the UZI domain are also conserved in these E3s and all their RING domains have large insertions (**Extended Data Fig. 6g, 7**). Hence, their E3 modules might also consist of a RING-UZI module like that experimentally confirmed with UBR4 **(Extended Data Fig. 6i**). Attempts to obtain soluble expression of the hemiRING lacking the UZI subdomain were unsuccessful, consistent with these folds being subdomains of the larger hemiRING-UZI module.

### The hemiRING is an E2 binding module

UBE2A variants cause UBE2A deficiency syndrome, an X-linked intellectual deficiency condition known as Nascimento type (MIM 300860), characterised by speech impairment, dysmorphic facial features and genital abnormalities ^51–53^. The atomic basis for the recognition of UBE2A by a cognate E3 and the pathogenic basis of these variants is unknown. When analyzed by size-exclusion chromatography a UBR4_xtal_ and UBE2A mixture eluted as a complex indicative of a stable interaction (**Fig. 4a**). To determine the structure of this complex we obtained crystals and collected diffraction data. A 3.2 Å structure was solved by molecular replacement using the apo structure obtained from UBR4_xtal_ and a UBE2A crystal structure (**Fig. 4b, c & Extended Data Fig. 8 9**) ^54^. As for the apo structure, the N-terminal ∼100 residues were also unresolved. The UBR4 molecule was equivalent to the apo structure (RMSD of 0.98 Å) except for the loop connecting αA and αB, gaining structural order, suggestive of E2 binding having a stabilizing effect on the hemiRING (**Fig. 4c**).

**Figure 4.**
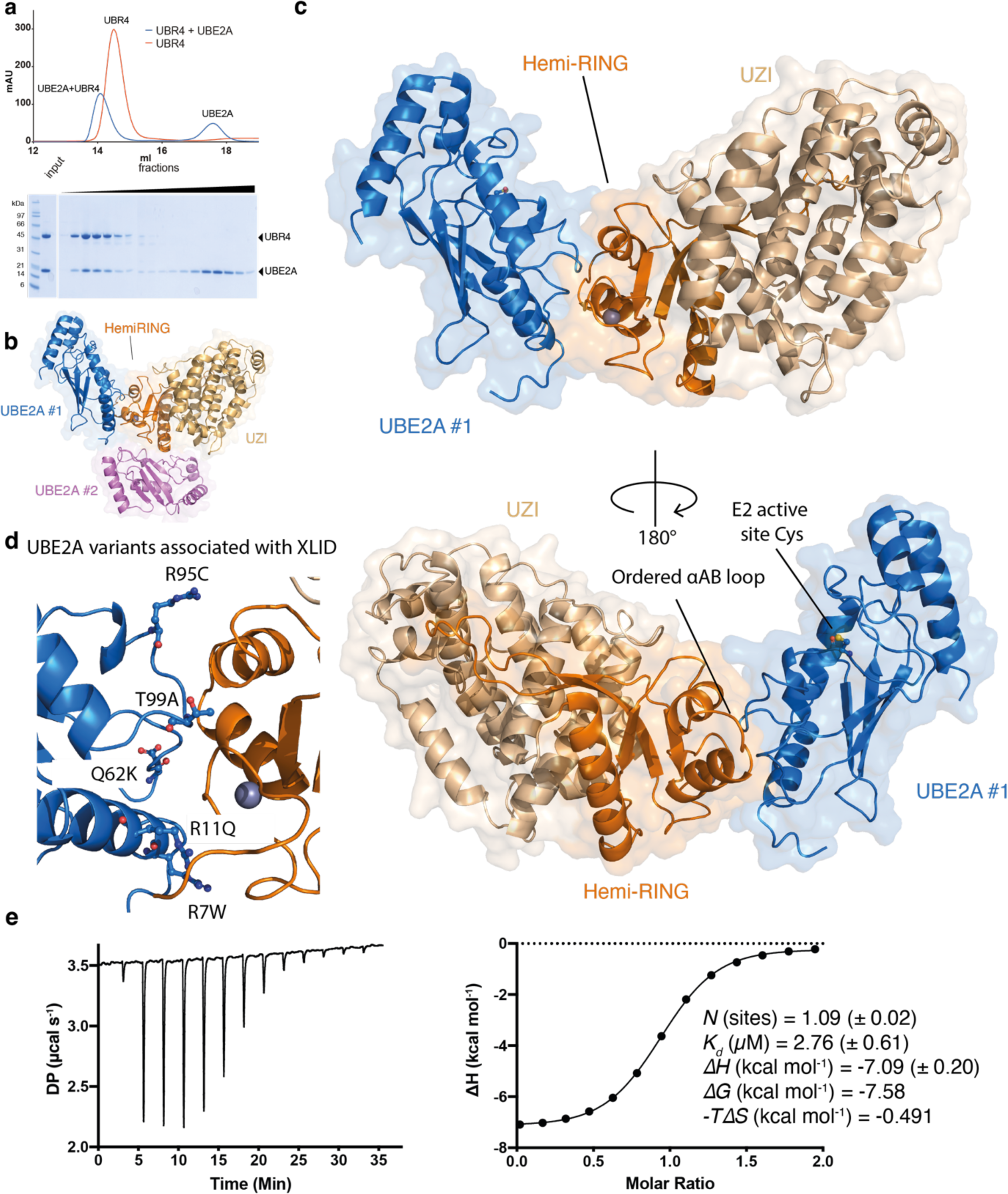
Crystal structure of UBR4 hemi-RING E3 in complex with E2 conjugating enzyme UBE2A. **a)** UBR4_xtal_ and UBE2A form a stable complex when analyzed by size-exclusion chromatography using a Superdex 200 10/300 GL (Cytiva Life Sciences) **b)** Asymmetric unit of the 3.2 Å crystallographic model for UBR4 hemiRING and UZI subdomains in complex with UBE2A. Structure is in in cartoon and transparent surface representation. Two UBE2A molecules are present in the asymmetric unit. Molecule UBE2A #1 is in blue and binds proximal to the zinc binding region of the UBR4 hemiRING. The second UBE2A molecule, UBE2A #2, is in violet. **c)** Enlarged views of the complex between UBR4 and UBE2A #1. A Cys88Lys UBE2A mutant was used for crystallization and this residue has been mutated to a cysteine *in silico*. **d)** X-linked intellectual disability (XLID) patient mutations cluster at the UBE2A-hemiRING interface. Residues are depicted in ball in stick and annotated with the pathogenic variant. **e)** ITC isotherm for UBE2A binding to UBR4_xtal_. A duplicate experiment was carried out with similar results (*N* = 0.904; *K_d_* = 2.58 μM; *11H* = -7.37 kcal mol^-1^).

Two UBE2A molecules are present in the asymmetric unit and the first E2 molecule (UBE2A #1) binds the hemiRING proximal to the zinc coordinating fold, similar to that observed with RING domains (**Fig. 4c**) ^48^. Several X-linked intellectual disability (XLID) patient mutations found in UBE2A cluster at this interface (**Figure. 4d**) ^53, 55, 56^. We could not establish any mechanistic significance for E2 molecule 2 (UBE2A #2) because mutation of E2 residues at this second interface had no clear effect on UBR4 autoubiquitination activity (**Fig. 4b, Extended Data Fig. 10**). However, in case the interaction was insensitive to our alanine scanning approach we also generated a steric UBR4 arginine mutant (Ser4930Arg), predicted to specifically clash with UBE2A #2, but this also demonstrated wild type levels of activity (**Extended Data Fig. 10**). Thus, the UBE2A #2 interaction is likely to be a consequence of crystallographic packing although in the context of endogenous protein it may be functional. We measured binding of UBE2A to UBR4_xtal_ by isothermal titration calorimetry (ITC) and determined that the free energy of binding (*ΔG*) is -7.58 kcal mol^-1^ (*K_d_* of 2.8 μM), with a minor entropic component (*TΔS =* 0.491 kcal mol^-1^) (**Fig. 4e**). Binding stoichiometry approximated unity indicating that if there were two E2 binding sites, the affinity of the second was below the affinity limit dictated by our ITC cell concentration of 74 μM.

### Interactions that govern E2-hemiRING interface are essential for E3 autoubiquitination

Although ancillary E3 elements that engage the backside of RAD6B (yeast orthologue of UBE2B) have been structurally delineated, the molecular basis for selective recognition of UBE2A/B by the core of an E3 module remains unknown^57^. The most notable interactions we observe arise from UBE2A residues Arg7, Arg8, Arg11, Arg95 and Ser97. Arg7, Arg8 and Arg11 reside in the UBE2A N-terminal α-helix whereas Ser97 and Arg95 are in loop 2. The guanidino groups of Arg7 and Arg11 hydrogen bond with side chain of UBR4 Glu4843 (**Fig. 5a**). Mutation of either of these residues to alanine, or mutating Glu4843 to arginine, severely impaired UBR autoubiquitination (**Fig. 5b**). These defects in activity are particularly insightful as Arg7Trp and Arg11Gln mutations are found in XLID patients and based on our findings, would impair UBE2A engagement by the UBR4 hemiRING, and in turn compromise substrate ubiquitination ^55, 56^. On the premise that UBR4 has neuronal functions, it is tempting to speculate that this observation represents the basis for XLID pathogenicity ^21, 23, 24^.

**Figure 5.**
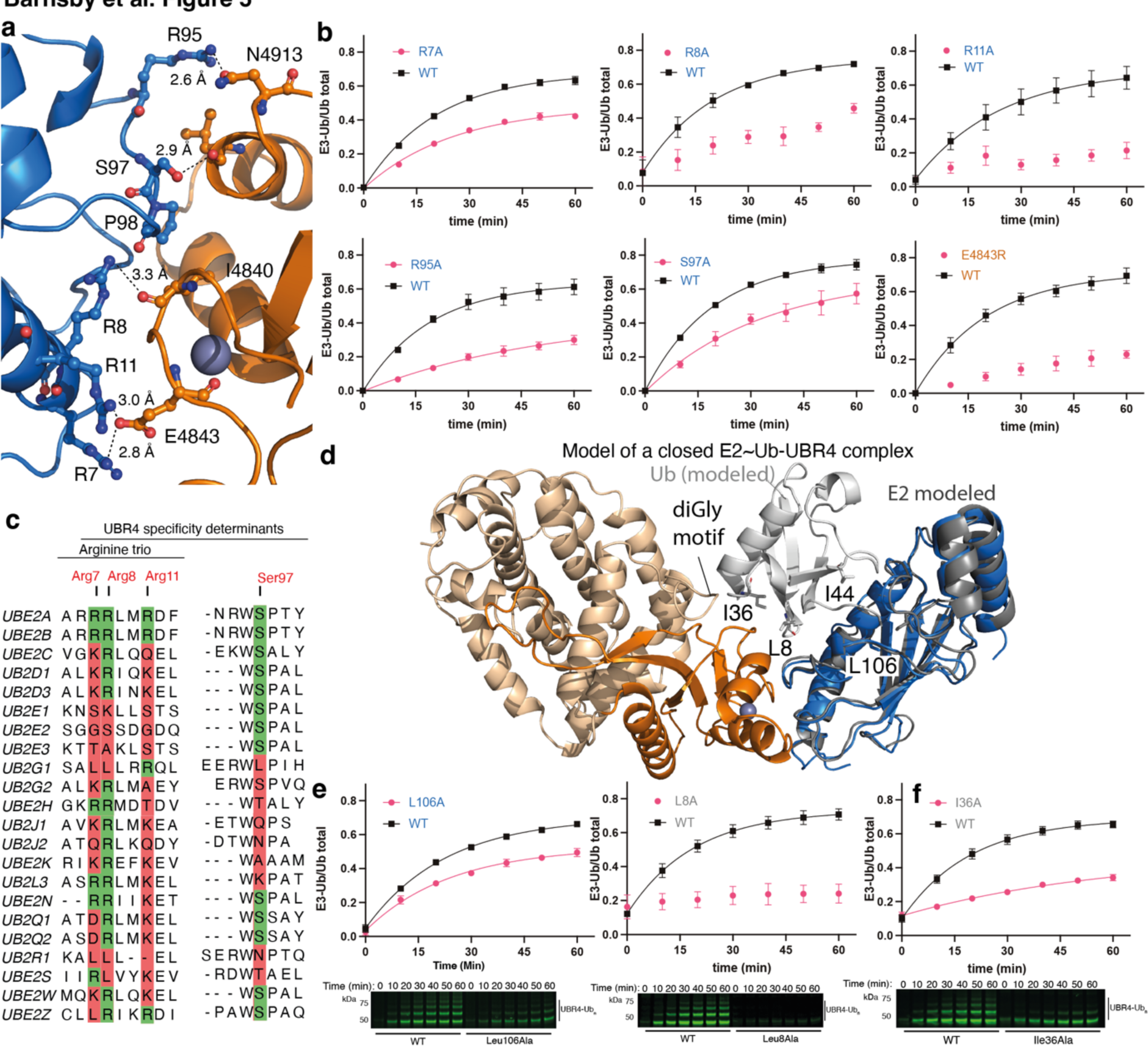
The UBE2A-UBR4 hemiRING interface and the requirement for a closed E2∼Ub conformation. **a)** Close up of the interface between UBE2A and the hemiRING. Four UBE2A arginine residues that are poorly conserved across mammalian E2s form key interactions with the hemiRING. UBE2A is in blue cartoon and key residues are in ball and stick. UBR4 hemiRING residues are in orange and key residues are in ball and stick. **b)** Mutational analysis of crystallographic interfacial UBE2A-hemiRING residues by in gel fluorescent autoubiquitination assay. Bars correspond to the standard error where n = 3. **c)** Sequence alignment performed with Jalview 2.11.2.5 using the Clustal algorithm for 22 mammalian E2 conjugating enzymes. Only UBE2A and UBE2B contain interface residues we experimentally found to be essential for optimal activity (Arg7, Arg8, Arg 11 and Ser97). **d)** Model of a ternary closed E2∼Ub-UBR4 complex obtained by superposition of UBE2D1∼Ub (PDB ID: 4AP4) onto the UBE2A E2 molecule (blue) in our experimental UBE2A-UBR4 structure. Ub residues essential for E3s that stabilize a canonical closed conformation are in grey ball and stick and are labelled. **e**) Mutation of Leu106 and Leu8 disrupts E3-dependent and independent lysine discharge because they make contacts at the E2-Ub interface of a close conformation. However, Leu106 in UBE2A is not solvent exposed and its mutation to alanine only partially impaired autoubiquitination. Bars correspond to the standard error where n = 3. **f)** Ile36 is not at the E2-Ub interface, and its mutation impairs activity with prototypical RING E3s that stabilize a closed E2∼ub conformation. Thus, it is likely to be only required for E3-dependent activity. Bars correspond to the standard error where n = 3.

UBE2A Ser97 also forms an important anchor point by hydrogen bonding through its side chain hydroxyl to the backbone carbonyl of UBR4 Ile4840, the side chain of which also makes hydrophobic contacts with UBE2A Pro98 (**Fig. 5a, b**). The only other interfacial contact of note is between UBE2A Arg95 and UBR4 Asn4913 and an Arg95Ala mutation significantly impaired autoubiquitination (**Fig. 5a, b**). This residue is also mutated in XLID patients (Arg95Cys) and would be similarly expected to compromise E3 activity^53^. With archetypal UBE2D1-3 isoforms glutamine (Gln92) is found in place of the arginine where its backbone carbonyl interacts with an arginine residue in canonical RING domains. This arginine within the RING domain acts as a “linchpin residue” that anchors the closed E2∼Ub conformation resulting in robust thioester activation ^9–11^. Structural and sequence analysis maps the equivalent site of this linchpin residue to Asn4913 in the hemiRING. Asparagine is also found in the cullin-associated RING protein Rbx1 where an alternative E2∼Ub activation mechanism is adopted^58^. However, this residue is unlikely to function as a linchpin in the conventional sense as it is too far from the E2 and might explain the inability of UBR4 to discernibly enhance discharge to free lysine.

To investigate whether the structurally resolved E2-UBR4 interface accounts for the exquisite specificity between UBR4 and UBE2A/B we carried out a sequence alignment of 22 E2s (**Fig. 5c**). A striking characteristic of UBE2A/B is the presence of a quartet of arginine residues (Arg7, Arg8, Arg11 and Arg95) and Ser97. Individual conservation of these residues varies from low to moderate, but the UBE2A/B α-helix 1 trio of arginine residues (Arg7, Arg8, Arg11), are unique to UBE2A/B. Thus, we propose that this more conservative structural motif is the minimal specificity determinant for UBE2A recognition by UBR4 and perhaps other E3s that are cognate for UBE2A/B.

### UBR4 activity involves a closed E2∼Ub conformation

Adapter-like E3s are understood to bind E2∼Ub and further stabilize its closed conformation thereby enhancing aminolysis ^9–11^. UBE2A can sample the closed conformation in the absence of an E3 and by modeling a ternary E2∼Ub-UBR4 complex we ascertained that this was sterically compatible with our complex structure (**Fig. 5d**) ^54^. To test the importance of a closed UBE2A∼Ub conformation, without assessment of whether UBR4 stabilizes it, we mutated residues that impede its formation and tested activity. With the UBE2D1-3 isoforms, Leu104 in the E2 crossover helix makes important hydrophobic contacts with the Ile44 patch of Ub and its mutation abolishes adapter-like E3 activity ^9–11, 59^. However, the equivalent residue in UBE2A is Leu106 which is positioned towards the core of the E2 fold and the exposed surface is more hydrophilic ^54^. Consistently, a Leu106Ala mutation only modestly impaired autoubiquitination activity (**Fig. 5e**). No detectable defect was observed in E3-independent lysine discharge indicating that the anomalous Leu104 site on UBE2A is a distinct requirement for UBR4-mediated autoubiquitination (**Extended Data Fig. 11**). Why this is the case is unclear. Another hydrophobic residue important for activity with RING exemplars studied to date is Ub Leu8, which packs against a distinct hydrophobic region of the E2 and the E3 ^9, 11^. An Leu8Ala mutant abolished autoubiquitination activity. However, E3-independent discharge to lysine was also significantly impaired, indicating a general defect in E2 activity (**Fig. 5e** **& Extended Data Fig. 11)**.

We next assessed whether UBR4_xtal_ stabilized the closed UBE2A∼Ub conformation, or whether it solely functioned as a scaffold that exerted its catalytic effect through substrate templating. Although UBR4_xtal_ did not enhance discharge to free lysine, this does not exclude a more tempered level of E2∼Ub activation that remained below the threshold of detection of untemplated discharge to free lysine. Indeed, an attenuated state of E2∼Ub activation is essential for tuning acceptor amino acid or substrate specificity ^40, 60^. A hydrophobic contact specific to E3 engagement is between Ub Ile36 and the distal RING protomer or with a non-RING priming element in monomeric RING E3s ^8, 9, 42, 61^. Interestingly, Ub Ile36 in our modelled ternary complex involving a closed UBE2A∼Ub conformation is proximal to a hydrophobic diGly motif in the UZI region (**Figure 5d**), suggesting the latter might serve as a non-RING priming element. We found that a Ub Ile36Ala mutant was defective in autoubiquitination whereas E3-independent discharge to free lysine was tolerant (**Fig. 5e** **& Extended Data Fig. 11**). These data suggest that the UZI subdomain acts as a tempered non-RING element by contacting Ile36 of Ub UBR4.

### The UBR4 E3 module is predicted to engage the backside of UBE2A

As a construct that approximated the crystallographically determined region of UBR4_xtal_ did not undergo autoubiquitination (UBR4_4828-5130_; **Fig. 1e**), we investigated the significance of the unresolved N-terminal region of UBR4_xtal_ (**Fig. 3b**). Initially, we sought insight into whether absence of autoubiquitination could be ascribed to the loss of modification sites. We excised the predominant autoubiquitination products from a Coomassie-stained SDS-PAGE gel and analyzed by data-dependent mass spectrometry **(Extended Data Fig. 12 & Extended Data File)**. This cursory assessment identified a single autoubiquitination site at Lys4814 (localization probability of 95 %), which is absent in UBR4_4828-5130_, suggestive of its removal being a potential cause for the loss of observable activity. Multiple isopeptide Ub linkages were also identified including Lys33, Lys48, Lys63, Lys11 and Lys6 (location probabilities of, 100 %, 100%, 96%, 85% and 83%, respectively). Thus, with the caveat that UBR4 coverage was incomplete, our observed oligomeric autoubiquitination pattern might consist of mixed linkage polyubiquitin extension from UBR4 residue Lys4814.

Despite the loss of a key autoubiquitination site accounting for the loss of activity with UBR4_4828-5130_, we were intrigued by the observation that E3 structural elements can bind non-canonically to the backside of E2s, including UBE2A, and potentiate or attenuate activity ^57, 62–64^. Furthermore, yeast UBR1 contains a helical element that binds in this manner ^4^. To explore whether the deleted region might also contribute to E3 activity we turned to the computational AlphaFold protein structure prediction tool and generated a model of a complex consisting of UBE2A and the unresolved residues in our crystal structures (**Fig. 6a**) ^50^. The model initiated with a kinked helix-helix followed by a 30-residue coil the mid-region of which facilitates extensive contacts with the UBE2A backside (**Fig. 6a**). The coil and this interaction were predicted with high confidence and following the coil is a short two stranded beta sheet ending with a 10-residue helix. Superposition of this model onto our E2-E3 crystal structure placed the C-terminal residue Ile4831 of UBR4_4730-4832_ 18 Å from Glu4832 in our experimentally determined E2-E3 complex (**Fig. 6b**). However, a contiguous polypeptide could be readily formed by rotation of the final 10-residue helix via the linker region between it and the short N-terminal beta sheet. Due to the experimental challenge in establishing whether the predicted helical extension might enhance E3 activity, as well as containing a primary ubiquitination site, we could not explore this intriguing possibility. However, in native context the helical region may well influence substrate ubiquitination. Collectively, our findings indicate that E3 modules can potentially be as large as ∼400 residues, which is of unprecedented scale for a single subunit system.

**Figure 6.**
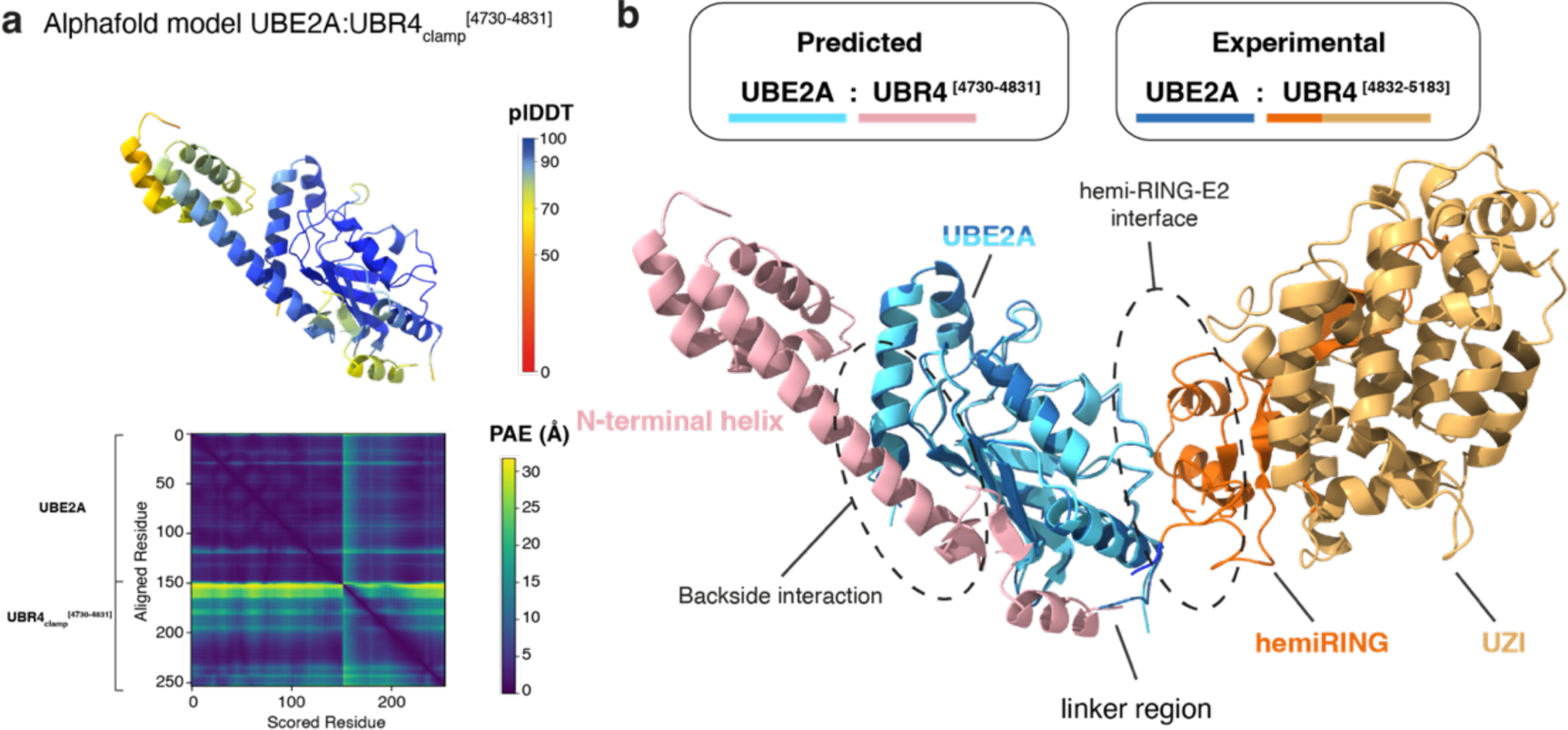
Computational identification and biochemical validation of an additional and essential component of the UBR4 E3 module. **a)** AlphaFold model of UBE2A in complex with the N-terminal region (residues 4730-4831) which is required for autoubiquitination but is unresolved in crystal structures. The protein is depicted in cartoon representation and colored by pIDDT score. The predicated alignment error (PAE) plot is also presented. **b)** E2 superposition of representative AlphaFold model with the experimental structure of the UBR4 hemiRING-UZI module.

## Discussion

Whilst UBR4 has been shown to destabilise N-degron substrates, the E3 module responsible has remained elusive. Herein we identify a novel adaptor-like E3 module within the giant N-degron E3 ligase UBR4. Whilst not discernible by primary sequence analysis, UBR4 contains an unusual zinc finger with partial structural similarity to the canonical cross-brace RING domain. However, only a single zinc ion is present and substituting for the second zinc ion is a hydrogen bonding network including a central water-mediated interaction. The fold is reminiscent of SP-RING domains found in SUMO E3 ligases of the Siz/PIAS family but the absence of the zinc ion distal to the E2 binding region has not been observed before. The loop region that coordinates the first zinc ion and a proximal α-helix present in canonical RING domains mediate interactions with E2 conjugating enzymes. Our crystal structure of a complex with a cognate E2 (UBE2A) reveals that this property is shared with the hemiRING. However, we identify important interfacial UBE2A and hemiRING residues providing insight into cognate E3 pairing with UBE2A/B. An arginine trio characteristic of UBE2A/B is central to its recognition by UBR4 as individual mutation impairs autoubiquitination activity. As mutation of these and other interfacial residues is found in X-linked intellectual disability patients, our structural provides insight into disease aetiology.

A somewhat unusual feature of the hemiRING is a pronounced loop. This loop, and perhaps the hemiRING itself, appear to require stabilization by an extreme C-terminal region composed of 11 alpha helices we refer to as the UBR Zinc-finger Interacting (UZI) domain, which is also mutated in disease. Unlike RING E3 exemplars, UBR4 does not discernibly enhance discharge to free lysine despite demonstrating robust E3 autoubiquitination activity. However, we find that the cognate E2 UBE2A has intrinsically high aminolysis activity and we propose that this compensates for the tempered nature of UBR4 catalysis. We reason that this intrinsically high state of UBE2A activity, presumably shared by UBE2B, explains why robust E3-mediated activation is unnecessary and accounts for the ability of these E2s to effect E3-independent targeted protein degradation^43^. Modelling of a closed E2∼Ub conjugate places Ub Ile36 in proximity of the UBR4 UZI domain and disruption of this putative interaction impairs autoubiquitination, leading us to assert that the UZI domain affects a tempered contribution to catalysis.

A structural feature of the UBR4 E3 module is an N-terminal helical region that was predicted to engage the backside of UBE2A by AlphaFold modelling. Binding of Ub to the backside of E2 enzymes enhances the processivity of polyUb chain formation ^63^. This event stabilizes E2 loop regions involved in RING binding and enhances the affinity of the interaction. Furthermore, helical elements from other E3s have been shown to engage the backside of UBE2A and modulate substrate ubiquitination. Thus, the predicted UBR4 helical region might similarly regulate E3 activity. Whether the N-terminal helical region further contributes to E3 selectivity for UBE2A/B remains to be tested.

The role of UBR4 E3 activity in promoting various diseases such as cancer and muscular atrophy would imply that modulation of UBR4 hemiRING E3 activity might have therapeutic value (e.g. by disruption of the UBE2A-hemiRING interface). As such, the mechanistic and structural insights obtained herein could be leveraged for the development of therapeutic modulators of UBR4-mediated substrate ubiquitination. Furthermore, although UBE2A/B have been shown to mediate efficient targeted protein degradation, they consist of a single core Ubc protein fold which has proven challenging to develop high affinity and selective ligands for ^65, 66^. The distinct structural features of the hemiRING-UZI module might provide a more tractable route to the development of ligands that indirectly recruit UBE2A allowing its intrinsically high lysine aminolysis activity to be exploited for therapeutic degrader applications.

Our findings uncover atomic resolution structure of a novel class of ubiquitin E3 ligase module and reveals the molecular basis for the selective recognition of UBE2A – an E2 that has not been structurally characterized in complex with a cognate E3. Our work demonstrates that novel adapter-like E3 modules remain to be identified and that they cannot be perceived by primary sequence analysis alone. Considering the discovery of unanticipated transthiolation E3s ^15, 67^, which only represent a small subset of the E3 ligase super family (< 10 %), it would seem probable that the scale and structural diversity of adapter-like E3s is also under appreciated.

## Acknowledgments

We also acknowledge MRC Reagents & Services for production of the recombinant E2 panel. We also thank Yogesh Kulathu for help with Alphafold calculations. This work was funded by the United Kingdom MRC (MC_UU_12016/8) and the Biotechnology and Biological Sciences Research Council (BB/P003982/1). L.B-G. was an awardee of an MRC Doctoral training partnership. We also acknowledge pharmaceutical companies supporting the Division of Signal Transduction Therapy (Boehringer-Ingelheim, GlaxoSmithKline, and Merck KGaA).

## Methods

### Expression and purification of UBR4 constructs

Glutathione-S-transferase (GST)-tagged UBR4 (WT and mutants) harboring a PreScission cleavage site were transformed into *E.coli* BL21(DE3) cells and grown overnight in an LB media starter culture supplemented with 200 µM zinc chloride and 100 μg/mL ampicillin at 37 °C with shaking. Starter culture was diluted 1:1000 into fresh LB supplemented with 200 µM zinc chloride and 100 μg/mL ampicillin and incubated at 37 °C until and OD_600_ of 0.8 was reached. Protein expression was induced with 0.3 mM isopropyl β-D-1-thiogalactopyranoside (IPTG) and cultures were incubated at 16 °C overnight.

Pellets were resuspended with buffer containing 20 mM HEPES-NaOH pH 7.4, 150 mM NaCl, 0.7 mM TCEP with 0.5 mg/mL lysozyme, 50 µg/mL DNase. Samples were sonicated on ice and clarified via centrifugation at 30,000 g for 45 minutes. Clarified lysates were incubated with GSH resin and washed via centrifugation with buffer containing 20 mM HEPES-NaOH pH 7.4, 150 mM NaCl, 0.7 mM TCEP. For elution, samples were incubated with 10 mM reduced glutathione for 10 minutes. For on resin tag cleavage, samples were incubated with C3 PreScission protease overnight at 4 °C. The eluted proteins were then further purified using size exclusion chromatography (HiLoad 16/600 Superdex 75 pg column or HiLoad 16/600 Superdex 200 pg column) using an ÄKTA Purifier FPLC system into buffer containing 20 mM HEPES-NaOH pH 7.4, 150 mM NaCl, 0.7 mM TCEP at 1 mL/min and collected in 1 mL fractions. Fractions of interest were then visualized via SDS-PAGE gel to assess purity and desired fractions were pooled and concentrated via spin concentrator, aliquoted and snap frozen prior to storage at -80 °C.

### Expression and purification of E2 conjugating enzymes

With exception of UBE2O, E2s were expressed in *E. coli* BL21(DE3) purified using glutathione Sepharose 4B or Ni-NTA resin, followed by size exclusion chromatography (SEC). N-terminal tags were cleaved with PreScission protease for E2s expressed from pGEX, pET156P and pET15b vectors whereas thrombin was used for E2s expressed from the pET28a vector (pET156P His-UBE2B, pET156P His-UBE2C, pET28a His-UBE2D1, pET28a His-UBE2D4, pET156P His-UBE2L3, pET28 His-UBE2S, pGEX6P-3-UBE2A and pGEX6P-1 UBE2R2). The N-terminal His tag on the remaining E2s was left in place and expressed from the following plasmids (pET28 His-UBE2D2, pET156P His-UBE2D3, pET156P His-UBE2E1, pET28a His-UBE2E2, pET28a His-UBE2E3, pET28a His-UBE2G1, pET28a His-UBE2G2, pET156P His-UBE2H, pET28a His-UBE2J1, pET28a His-UBE2J2, pET156P His-UBE2K, pET15b His-UBE2N, pET28a His-UBE2Q1, pET15b His-UBE2Q2, pET28a His-UBE2R1, pET15b6P His-UBE2T, pET28a His-UBE2V1, pET15b His-UBE2V2, pET28a His-UBE2W and pET15b His-UBE2Z). UBE2O was expressed in Sf9 insect cells. Protein was purified using Ni-NTA affinity followed by SEC and the His tag was left in place.

### Autoubiquitination assays

Assays were made up from a 10X buffer (400 mM Na_2_HPO_4_, 1.5 M NaCl, 10 mM TCEP) containing final concentrations of 40 mM Na_2_H_2_PO_4_ pH 8, 150 mM NaCl, 5 mM MgCl_2_, 1 mM TCEP, ATP (5 mM), cleaved UBR4 construct (3 µM), E2 (5 µM), E1 (0.5 µM), ubiquitin (50 µM). Reactions were incubated at 37 °C for 30 minutes and quenched by dilution with 4X LDS buffer containing 680 mM 2-mercaptoethanol and resolved via SDS-PAGE gel, visualised by either Coomassie staining or western blotting.

For full-length UBR4 auto-ubiquitination assays, HEK293 cells stably overexpressing HA-UBR4 were lysed in lysis buffer (50 mM Tris-HCl pH 7.5, 1 mM EGTA, 1 mM EDTA, 10 mM glycerophosphate, 50 mM sodium fluoride, 5 mM sodium pyrophosphate, 1 mM sodium vanadate, 0.27 M sucrose, 1% NP-40, 0.2 mM PMSF, 1 mM benzamidine, 1 mM TCEP) supplemented with complete EDTA-free protease inhibitor cocktail (Roche 11873580001). Lysate was then centrifuged for 10 min at 16,200 x g, and the supernatant was collected. Full length HA-UBR4 was immunoprecipitated using anti-HA sepharose resin for one hour at 23 °C, followed by a wash in phosphate buffer (50 mM sodium phosphate pH 7.5, 150 mM NaCl, 1 mM TCEP). For the E2 panel, the resin was combined with reaction mix containing UBE1 (500 nM), ubiquitin (5 µM), ATP (10 mM) and various E2s (5 µM for UBE2E3, G1, J1, J2, R2, S and 1.5 µM for UBE2O) in 50 mM sodium phosphate pH 7.5, 150 mM NaCl, 5 mM MgCl_2_, 0.75 mM TCEP. Concentrations of 12.5 µM ubiquitin and 1 mM TCEP were used for the full-length UBR4 mutant’s experiment. Reactions were performed at 37°C for one hour and stopped by the addition of LDS loading buffer containing β-mercaptoethanol (Invitrogen NP0007). Samples were heated for 10 min at 70 °C and loaded on 4-12% Bis-Tris (Invitrogen NP0323) and 3-8% Tris-Acetate (Invitrogen EA03785) polyacrylamide gels for Coomassie stain (Instant Blue Abcam AB119211) or Western blotting, respectively.

### Crystallization of UBR4_xtal_ and UBR4_xtal_-UBE2A complex

Initially, commercially available crystallisation conditions were screened in 96-well format. Proteins for crystal screening were expressed and purified as above for cleaved protein. Plates were set up with the Mosquito Crystal and Dragonfly liquid handling robots (SPT Labtech). Plates were then covered and left at either 4 °C or room temperature and imaged periodically. Conditions successful at yielding crystals (10 mM Na_2_HPO_4_, pH 6.5, 13% PEG20000, 22°C and 4 °C) were replicated on a larger scale in 24 well plates which were incubated at room temperature. 1 mL total volume of each condition was placed in each well and covered with a glass cover slip. Each slip contained a 2 µL hanging drop (1 µL protein, 1 µL buffer condition) with protein concentration at 10 mg/mL (5 mg/mL final per drop). For the UBR4_xtal_-UBE2A complex, the 6xHis tag was cleaved from a UBE2A Cys88Lys mutant (DU 65350) with Tobacco Etch Virus (TEV) protease was mixed with an equimolar amount of cleaved GST PreScission UBR4_xtal_ (DU 65064) and crystals were obtained in hanging drops 0.1 M Bis-Tris, pH 6.4, 15 % PEG10000, 0.2 M Ammonium acetate.

Crystals were collected and cryo-protected with the well condition supplemented with 25% ethylene glycol followed by plunge vitrification in liquid nitrogen. Crystals were screened at Diamond Light Source, Oxford UK beam lines I24 (UBR4_xtal_) or I04-1 (UBE2A-UBR4_xtal_) via remote collection. For UBR4_xtal_ crystals an X-ray fluorescence scan at the zinc K-absorption edge was performed. Based on the scan the peak wavelength was chosen as λ = 9671.0 eV (1.2820Å) and the inflection point wavelength as λ = 9663.0eV (1.2831Å). Data were collected at the zinc edge to allow measurement of the anomalous signal for phasing. The data were indexed, integrated, and scaled using DIALS^68^ and phased using CRANK2 (ver. 2.0.1). Refinement was carried out with Phenix (ver. 1.17.1) ^69^, between rounds of refinement models were manually improved using Coot ^70^ and Final Ramachandran statistics were favored 98.54 %, allowed 1.46 % and outliers 0.00 %. Clash Score was 3.54.

For the UBE2A-UBR4_xtal_ complex data was collected at 0.9118 Å and was indexed and integrated using DIALS. Data were scaled and merged using Aimless (0.7.4). Phasing was achieved by molecular replacement using Phaser (ver. 2.8.3), the UBR4_xtal_ structure and UBE2A (PDB 6CYO) ^54^ were used as search models. Refinement was carried out with Phenix (ver. 1.17.1) ^69^, between rounds of refinement models were manually improved using Coot (ver. 0.9.5) ^70^. Final Ramachandran statistics were favoured 95.70 %, allowed 4.30 % and outliers 0.00 %. Clash Score was 4.42.

### Preparation of Cy3b labelled ubiquitin

Ubiquitin with an N-terminal His-tag followed by a cysteine residue for Cy3b conjugation and a TEV (Tobacco-etch virus) cleavage site (DU 29939) was expressed in BL21 cells as described above and purified using Ni-affinity chromatography. The His-tag was cleaved with TEV protease as described above and protein was buffer exchanged into degassed buffer containing 50 mM HEPES-NaOH, pH 7.5, 0.5 mM TCEP prior to dye conjugation. Protein was concentrated to 2 mg/mL and 200 µL was mixed with Cy3b-maleimide for a final concentration of 150 nM in 300 µL. Protein was then incubated at 25 °C for 2 hours with agitation. Reaction was monitored by LC-MS (Agilent 1200 HPLC, 6130 Single Quad) then purified using a P2 Centri-Pure desalting column using the same degassed buffer. Concentration was then checked, and the protein was aliquoted, snap frozen and stored at - 80 °C.

### UBR4 autoubiquitination under single turnover E2∼Ub discharge conditions

6xHis cleaved wild-type or mutant E2s (10 µM) were charged with ubiquitin labelled with Cy3b at a concentration of 12.5 µM, in a buffer of 20 mM Na_2_H_2_PO_4_ pH 8, 150 mM NaCl, 1 mM TCEP, along with 0.5 µM E1 and 5 mM ATP for 20 minutes at 37°C. Samples were cooled on ice for 2 minutes followed by addition of pan E1 inhibitor Compound 1 (25 µM) as previously described ^15^, and incubated for 10 min at room temperature. This was then mixed with another sample containing either UBR4 (WT or mutant) 5 µM, or buffer in the case of no E3 controls. Samples were taken at indicated time points and quenched with 4X LDS loading dye (ThermoFisher). Gels were imaged via ChemiDoc, and data analysed via ImageJ and Prism.

### Lysine discharge assay

WT or mutant E2s (10 μM) were charged with labelled ubiquitin (Cy3/Cy3b/Cy5) (12.5 μM) as above. An equal volume of sample containing 20 mM Na_2_PO_4_ ∼pH 7.5, 20 mM L-Lysine, or buffer alone for control reactions, was added and a time point taken at t=0. Further samples were taken at the indicated time points and quenched with 4X LDS loading buffer (ThermoFisher).

### Transient transfection of wild type and mutant full length UBR4

HEK293 cells were grown in 100 mm dishes, and transfected with wild type, Cys4890Ala, or His4887Ala N-terminal HA-tagged UBR4 coding plasmids (DU 71005 and 71006), or the empty vector, using Lipofectamine 2000 (Invitrogen 11668-019). Briefly, plasmid DNA (10 µg) and Lipofectamine 2000 (25 µg) were diluted in Opti-MEM (GIBCO 31985-062), combined, and incubated at room temperature for 20 min before adding onto cells. Cells were maintained in DMEM (GIBCO 11960-085) 10% fetal bovine serum (Sigma F7524), 100 U/mL penicillin-streptomycin (GIBCO 15140122), 2 mM L-glutamine (GIBCO 25030024) at 37 °C in 5% CO_2_, and a period of 24 h post-transfection was observed before harvesting to allow protein expression.

### Stable UBR4 expressing cell line

Full-length UBR4 stable cell lines were created by co-transfecting untagged (DU65532) or N-terminal HA-tagged UBR4 (DU65964) coding plasmids (7.5 µg) and pOG44 (2.5 µg) into HEK293 Flp-In T-REx cells. Selection and maintenance of cells that underwent recombination started 24 h later by including hygromycin (50 µg/ml, Invivogen ant-hg-5) in the culture media. Induction of UBR4 expression was achieved by supplementing media with tetracycline (1 µg/ml, Sigma T7660).

### Western blotting

Electrophoresis was performed at 200 V and transferred on PVDF membrane using a Tris-glycine buffer (48 mM Tris, 39 mM glycine, 20% methanol) at 95 V for 3 h. Membranes were incubated one hour in 5% milk before adding the primary antibody (anti-HA 3F10, Roche 27573500, 1:2500, anti-Ubiquitin P4D1, Biolegend, 1:10,000, anti-UBR4/p600 ab86738, Abcam, 1:5000, anti-FLAG M2, Sigma F1804, 1:5000, anti-Vinculin ab129002, Abcam, 1:10,000). After rinsing, secondary antibody was added (anti-rat Cell Signaling 7077S, 1:5000, anti-mouse Cell Signaling, 7076S, 1:5000, anti-rabbit IRDye 680RD, LI-COR 926-68071, 1:20,000, anti-mouse IRDye 800CW, LI-COR 926-32210, 1:20,000). Following further washing, some membranes were incubated in chemiluminescent substrate (ECL Pierce 32106). Chemiluminescence was captured by radiographic films or an electronic imaging system (ChemiDoc MP Bio-Rad) running Imagelab Touch Software (ver. 2.3.0.07). Near-infrared signal was assessed using a LI-COR Odyssey CLx instrument running Image Studio (ver. 5.2).

### Isothermal Titration Calorimetry (ITC)

ITC experiments were carried out using a MicroCal PEAQ-ITC instrument (Malvern Panalytical). UBR4 constructs were placed in the cell (74 μM) and UBE2A was placed in the syringe at 10-fold higher concentration. Experiments utilised an initial injection (0.4 µL) followed by 13 injections (3 µL). A reference response of UBE2A titrated into buffer was subtracted using the manufacturer software routine to account for the dilution enthalpy of the titrant. All experiments were carried out at 25 °C and the data generated were analysed using manufacturer software and PRISM (Graphpad) for figure generation.

### Ubiquitination Site Mapping by Mass Spectrometry LC-MS/MS Analysis

Peptides generated by trypsin treatment of the excised gel slice were resuspended in 5 % formic acid in water and injected on an UltiMate 3000 RSLCnano System coupled to an Orbitrap Fusion Lumos Tribrid Mass Spectrometer (Thermo Fisher Scientific). Peptides were loaded on an Acclaim Pepmap trap column (Thermo Fisher Scientific #164750) with prior analysis on a PepMap RSLC C18 analytical column (Thermo Fisher Scientific #ES903) and eluted on a 120 min linear gradient from 3 to 35% Buffer B (Buffer A: 0.1% formic acid in water, Buffer B: 0.08% formic acid in 80:20 acetonitrile:water (v:v)). Eluted peptides were then analysed by the mass spectrometer operating in data dependant acquisition mode. Peptides were searched against a reduced database containing only the four proteins used in this assay (ubiquitin, His-UBA1, UBE2A and UBR4) using MaxQuant (v2.1.3.1) 71. All parameters were left as default except for the addition of Deamidation (N, Q) and GlyGly (Protein N-term, K, C, S, T, Y) as variable modifications modification and with the PSM, Protein and Site FDR increase to 1.00. MS/MS Spectra of interesting GlyGly peptides were manually inspected.

