## Supplemental Figures for "An Atypical E3 Ligase Module in UBR4 Mediates Destabilization of N-degron Substrates"

**EXTENDED DATA FIGURES**

Lucy Barnsby-Greer[1,†], Peter D. Mabbitt[1,2†], Marc-Andre Dery[1], Daniel R. Squair[1], Nicola T. Wood[1], Sven Lange[1] & Satpal Virdee[1]*

MRC Protein Phosphorylation and Ubiquitylation Unit, University of Dundee, Scotland, UK, DD1 5EH

[1] MRC Protein Phosphorylation and Ubiquitylation Unit, University of Dundee, Scotland, UK, DD1 5EH

[2]Scion, Titokorangi Drive, Private Bag 3020, Rotorua 3046, New Zealand

[†]These authors contributed equally

*Corresponding author

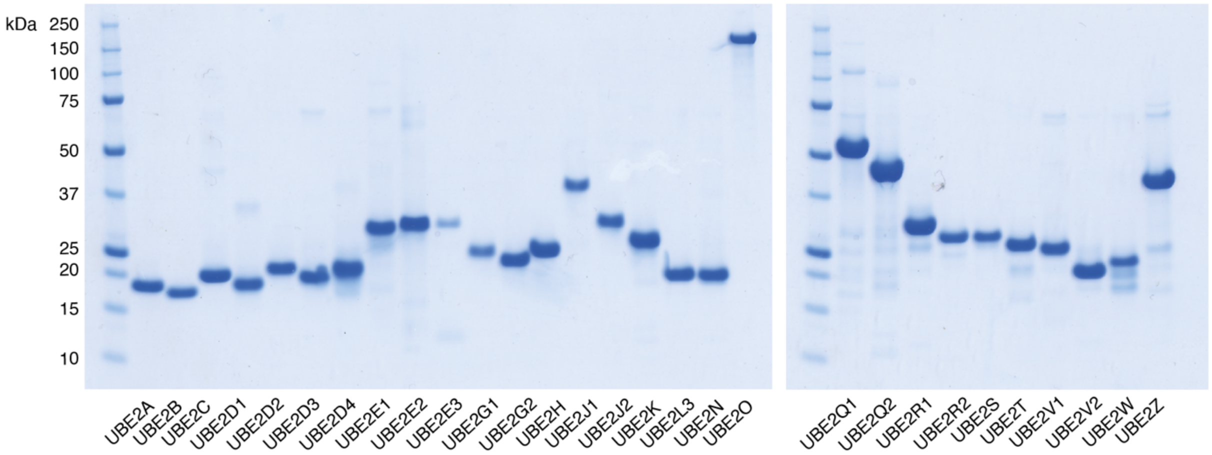

**Extended Data Figure 1. Recombinant panel of E2 conjugating enzymes used in UBR4 activity assay.** E2s were expressed in *E. coli* and purified by affinity chromatography using established methods. UBE2A, UBE2B, UBE2C, UBE2D1, UBE2D4, UBE2L3, UBE2R2 and UBE2S were untagged whereas the remaining E2s bared N-terminal His tags.

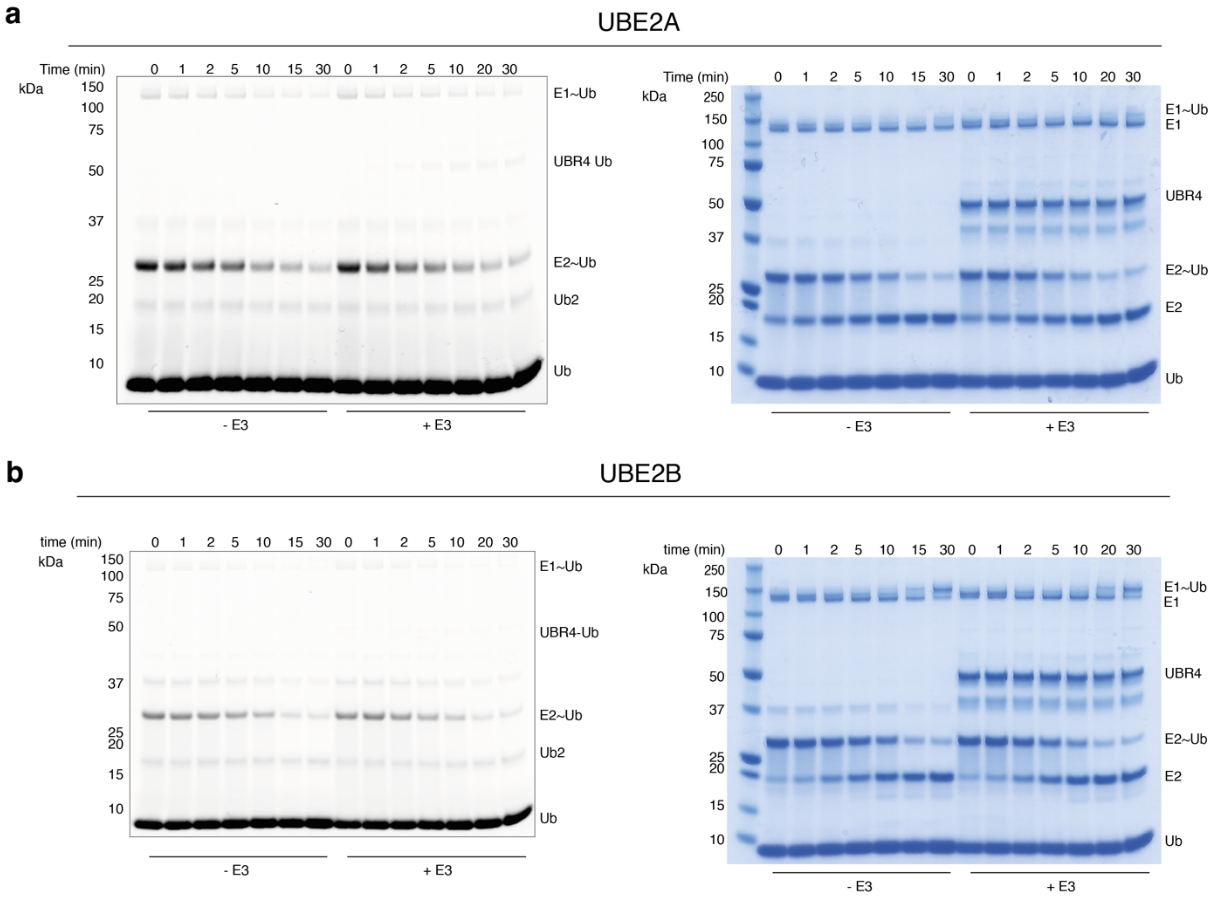

**Extended Data Figure 2. Assessment of UBR4-dependent discharge from UBE2A and UBE2B to free lysine. a)** Single turnover discharge of Cy3b-labelled Ub from UBE2A (5 μM) to lysine in the presence and absence of UBR4$_{xtal}$ (5 $\mu$M). Gels were visualized by in-gel fluorescence (left) and by Coomassie staining (right). **b)** As above but for UBE2B. Both of these experiments with UBR4 at 5 $\mu$M were carried out once.

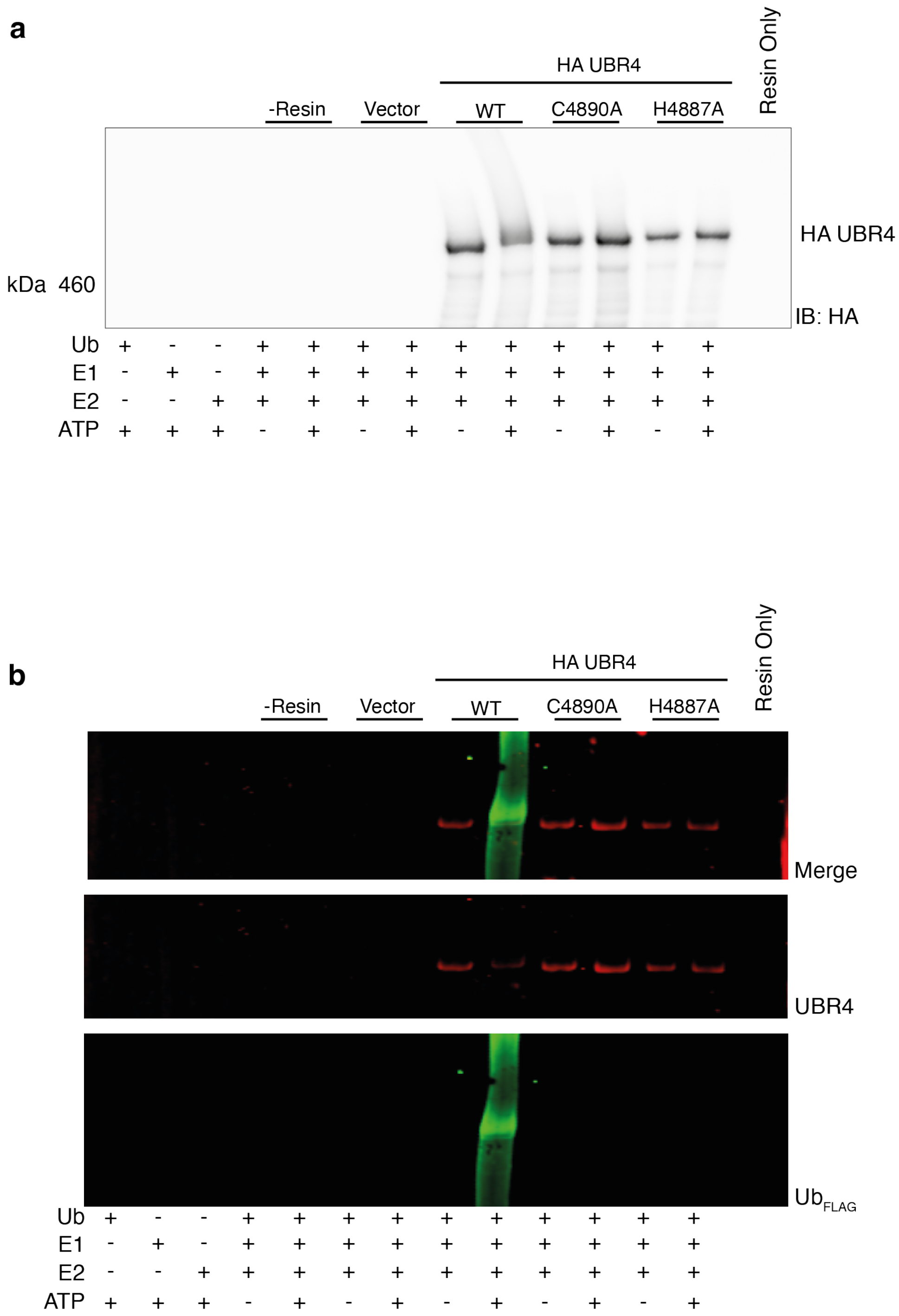

**Extended Data Figure 3. Autoubiquitination assay for full-length UBR4 C4890A and H4887A mutants. a)** Wild type and the corresponding HA-tagged UBR4 mutants were transiently overexpressed in HEK293 cells and immunoprecipitated against HA-sepharose resin. Washed resin was combined with E1 (500 nM), FLAG-ubiquitin (5 µM), ATP (10 mM) and UBE2A E2 (5 µM). Reactions were incubated at 37 °C for one hour, stopped by the addition of reducing LDS loading buffer and visualized by anti-HA immunoblot. **b)** Near-infrared imaging of the membranes was also carried out with a LI-COR Odyssey CLx system. For the UBR4 and FLAG channels anti-rabbit IRDye 680RD and anti-mouse IRDye 800CW were used, respectively.

| | $UBR4_{xtal}$ |
|---|---|
| **Data collection** | |
| Space group | I 2 2 2 |
| Cell dimensions | |
| *a*, *b*, *c* (Å) | 67.131 84.644 148.13 |
| $\alpha, \beta, \gamma$ (°) | 90 90 90 |
| Resolution (Å) | 74.07 - 1.8 (1.864 - 1.8) |
| $R_{merge}$ | 0.02395 (0.2907) |
| $I / \sigma I$ | 23.84 (2.56) |
| CC1/2 | 0.999 (0.872) |
| Completeness (%) | 91.33 (91.67) |
| Redundancy | 2.0 (2.0) |
| **Refinement** | |
| Resolution (Å) | 74.07 - 1.8 |
| No. reflections | 71,900 (6,996) |
| $R_{work}$ / $R_{free}$ | 0.1981/ 0.2220 |
| No. atoms | |
| Protein | 2,723 |
| Ligand/ion | 9 |
| Water | 408 |
| *B*-factors | |
| Protein | 26.84 |
| Ligand/ion | 38.66 |
| Water | 36.44 |
| R.m.s. deviations | |
| Bond lengths (Å) | 0.003 |
| Bond angles (°) | 0.56 |

*Values in parentheses are for highest-resolution shell.

**Extended Data Figure 4. Data collection and refinement statistics for $UBR4_{xtal}$.** Data were collected from a single crystal using single-wavelength anomalous diffraction (SAD).

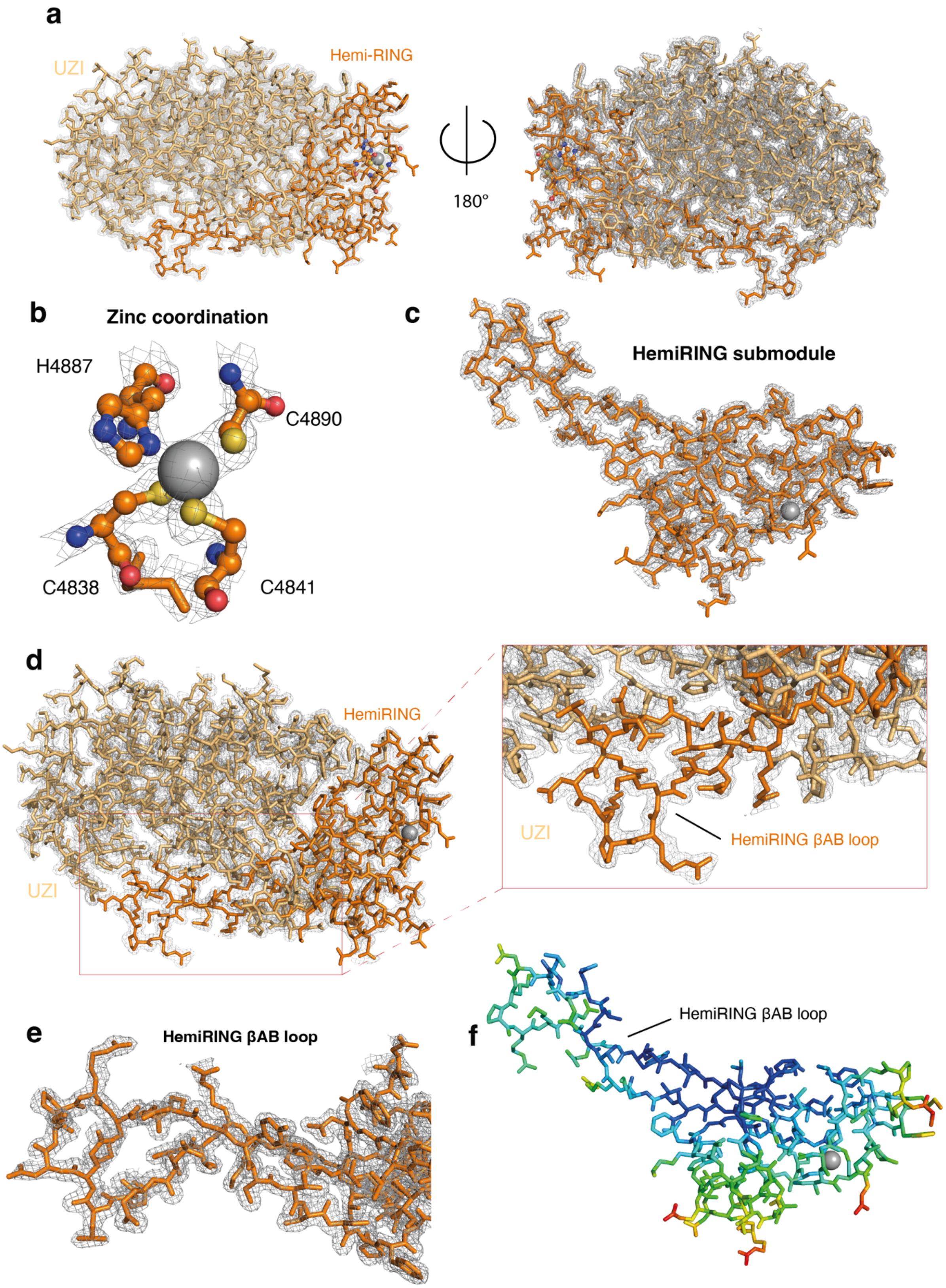

**Extended Data Figure 5. Representative views of the crystallographic model of the UBR4 E3 module. a)** A single molecule, corresponding to residues 4831-5183, was found in the asymmetric unit. The model is represented in stick where the hemiRING submodule is coloured orange and the UBR Zinc finger-Interacting (UZI) subdomain is coloured wheat. **b)**

Close up of the coordination network towards the single $Zn^{2+}$ ion in the hemiRING submodule. Mutation of these residues ablates E3 ligase activity. **c)** Isolated view of the hemiRING submodule. Inserted into the hemiRING is a pronounced extension made up of the βAB loop. **d)** Interaction between the hemiRING βAB loop and the UZI subdomain, the latter appearing to be integral to the stability of the βAB loop. **e)** Close up isolated view of the hemiRING βAB loop. For panels a-e the mesh corresponds to a $2|F_{obs}|-|F_{calc}|$ electron density map contoured at 1.0 σ. **f)** HemiRING sub-module colored by B-factor (blue is lowest and red is highest).

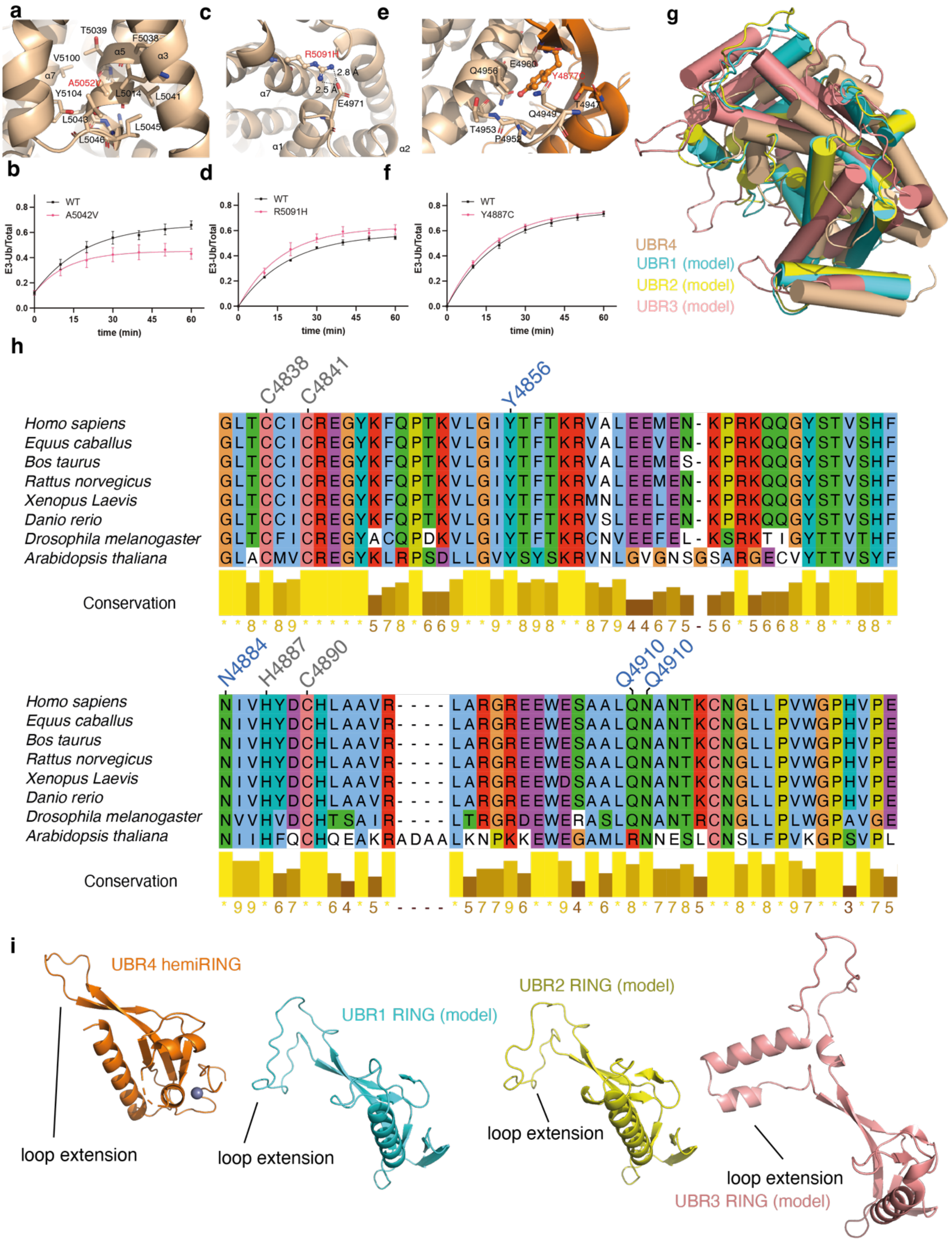

a
T5039
F5038
V5100
α5
α3
α7
A5042V
Y5104
L5014
L5041
L5043
L5045
L5046
c
R5091H
2.8 Å
2.5 Å
α7
E4971
α1
α2
e
Q4956
E4960
Y4887C
T4947
Q4949
T4953
P4952
b
WT
A5042V
E3-Ub/Total
time (min)
d
WT
R5091H
E3-Ub/Total
time (min)
f
WT
Y4887C
E3-Ub/Total
time (min)
g
UBR4
UBR1 (model)
UBR2 (model)
UBR3 (model)
h
C4838
C4841
Y4856
Homo sapiens GLTCCICREGYKFQPTKVLGIYTFTKRVALEEMEN-KPRKQQGYSTVSHF
Equus caballus GLTCCICREGYKFQPTKVLGIYTFTKRVALEEVEN-KPRKQQGYSTVSHF
Bos taurus GLTCCICREGYKFQPTKVLGIYTFTKRVALEEMES-KPRKQQGYSTVSHF
Rattus norvegicus GLTCCICREGYKFQPTKVLGIYTFTKRVALEEMEN-KPRKQQGYSTVSHF
Xenopus Laevis GLTCCICREGYKFQPTKVLGIYTFTKRMNLEELEN-KPRKQQGYSTVSHF
Danio rerio GLTCCICREGYKFQPTKVLGIYTFTKRVSLEEFEN-KPRKQQGYSTVSHF
Drosophila melanogaster GLTCFICREGYACQPDKVLGIYTFTKRCNVEEFEL-KSRKTIGYTTVTHF
Arabidopsis thaliana GLACMVCREGYKLRPSDLLGVYSYSKRVNLGVGNSGSARGECVYTTVSYF
Conservation
**8*89*****578*669**9*898**879446 75-56*5668*8**88*
N4884
H4887
C4890
Q4910
Q4910
Homo sapiens NIVHYDCHLAAVR----LARGREEWESAALQNANTKCNGLLPVWGPHVPE
Equus caballus NIVHYDCHLAAVR----LARGREEWESAALQNANTKCNGLLPVWGPHVPE
Bos taurus NIVHYDCHLAAVR----LARGREEWESAALQNANTKCNGLLPVWGPHVPE
Rattus norvegicus NIVHYDCHLAAVR----LARGREEWESAALQNANTKCNGLLPVWGPHVPE
Xenopus Laevis NIVHYDCHLAAVR----LARGREEWDSAALQNANTKCNGLLPVWGPHVPE
Danio rerio NIVHYDCHLAAVR----LARGREEWESAALQNANTKCNGLLPVWGPHVPE
Drosophila melanogaster NVVHVDCHTSAIR----LTRGRDEWERASLQNANTRCNGLLPLWGPAVGE
Arabidopsis thaliana NIIHFQCHQEAKRADAALKNPKKEWEGAMLRNNESLCNSLFPVKGPSVPL
Conservation
*99*67**64*5*----*57796**94*6*8*7785**8*8*97**3*75
i
UBR4 hemiRING
loop extension
UBR1 RING (model)
loop extension
UBR2 RING (model)
loop extension
loop extension
UBR3 RING (model)

**Extended Data Figure 6. Episodic ataxia (EA) patient mutations within the hemiRING-UZI module, structural superposition of UBR1-4 UZI subdomains and AlphaFold models of RING domains from UBR1, UBR2 and UBR3. a)** Structural context residue Ala5042. An Ala5042Val mutation is associated with EA patients. **b)** Introduction of the Ala5042Val mutation impaired UBR4 autoubiquitination ($n$ = 3 and bars represent the standard error). **c)** Structural context of R5091 which is mutated to histidine in EA patients. **d)** The Arg5091His mutant had slightly reduced autoubiquitination activity ($n$ = 3 and bars represent the standard error). **e)** Structural context of Tyr4877Cys which is mutated to cysteine in EA patients. **f)** The Tyr4877Cys mutation demonstrated a modest defect in UBR4 autoubiquitination ($n$ = 3 and bars represent the standard error). **g)** Structural superposition of AlphaFold models of UZI subdomains predicted to exist within *Homo sapiens* UBR1-3 with our experimentally determined structure of the UZI subdomain from UBR4. **h)** The human UBR4 hemiRING was aligned against selected orthologues. All residues involved in coordination of the single zinc ion are conserved and are labelled in grey. Except for Gln4910, which is an arginine residue in *Arabidopsis*, all residues that form the hydrogen bonding network that replaces the canonical distal zinc coordination site are also conserved and labelled in blue. However, the Gln4910 side chain is solvent exposed suggestive of side chain tolerance at this position. Alignment and figure generation was carried out with Jalview 2.11.2.5 using the Clustal algorithm. **i)** AlphaFold models of the canonical RING domains from UBR1-3 indicates that like the UBR4 hemiRING, they contain pronounced insertions within the RING domain that also interact with their respective UZI domain that might also result in stabilization of the former.

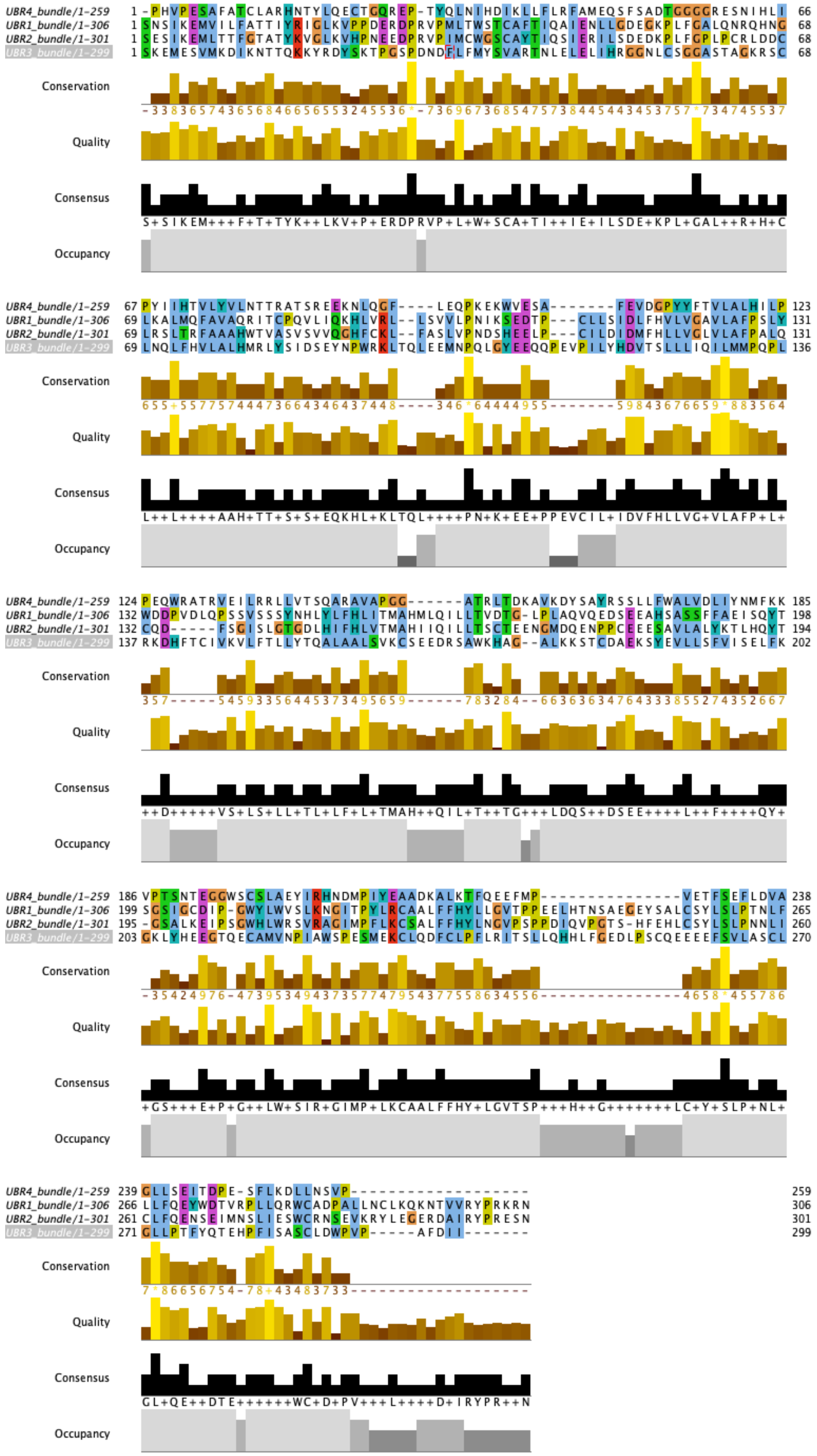
UBR4_bundle/1-259
UBR1_bundle/1-306
UBR2_bundle/1-301
UBR3_bundle/1-299
Conservation
Quality
Consensus
Occupancy

**Extended Data Figure 7. Sequence alignment of UBR1-4 UZI subdomains.** Sequences were aligned with Clustal using Jalview 2.11.2.5.

| | UBR4-UBE2A complex |
|---|---|
| **Data collection** | |
| Space group | P 43 21 2 |
| Cell dimensions | |
| *a*, *b*, *c* (Å) | 89.1569 89.1569 263.306 |
| α, β, γ (°) | 90 90 90 |
| Resolution (Å) | 65.83 - 3.2 (3.315 - 3.2) |
| $R_{merge}$ | 0.2103 (1.251) |
| $I / \sigma I$ | 8.04 (1.35) |
| CC1/2 | 0.999 (0.983) |
| Completeness (%) | 99.62 (99.50) |
| Redundancy | 12.5 (13.1) |
| **Refinement** | |
| Resolution (Å) | 65.83 - 3.2 |
| No. reflections | 229,999 (23,721) |
| $R_{work}$ / $R_{free}$ | 0.2282/ 0.2623 |
| No. atoms | |
| Protein | 5,186 |
| Ligand/ion | 1 |
| Water | 0 |
| *B*-factors | |
| Protein | 82.05 |
| Ligand/ion | 63.83 |
| Water | N/A |
| R.M.S. deviations | |
| Bond lengths (Å) | 0.002 |
| Bond angles (°) | 0.45 |

*Values in parentheses are for highest-resolution shell.

**Extended Data Figure 8. Data collection and refinement statistics for UBE2A-UBR4$_{xtal}$.** Data were collected from a single crystal and phased by molecular replacement.

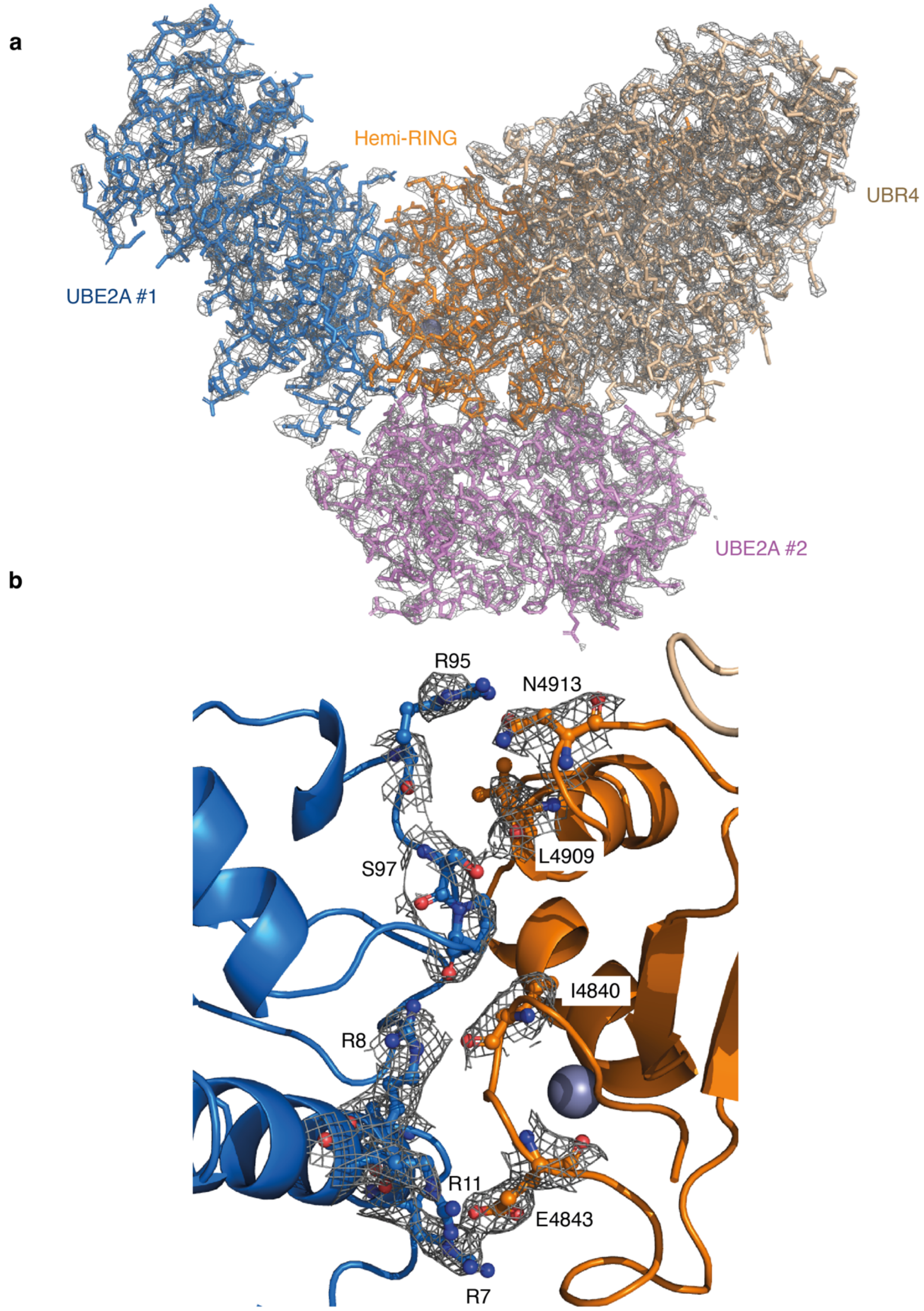

**Extended Data Figure 9.** ***2|$F_o$|-|$F_c$| electron density map for the UBE2A-UBR4 complex. a)*** The Complete asymmetric unit in stick format is overlaid with the 2|$F_o$|-|$F_c$| map**. b)** The UBE2A-hemiRING interface. Key residues are depicted in ball and stick and their corresponding 2|$F_o$|-|$F_c$| map is overlaid. Maps contoured at 1.0 $\sigma$**.**

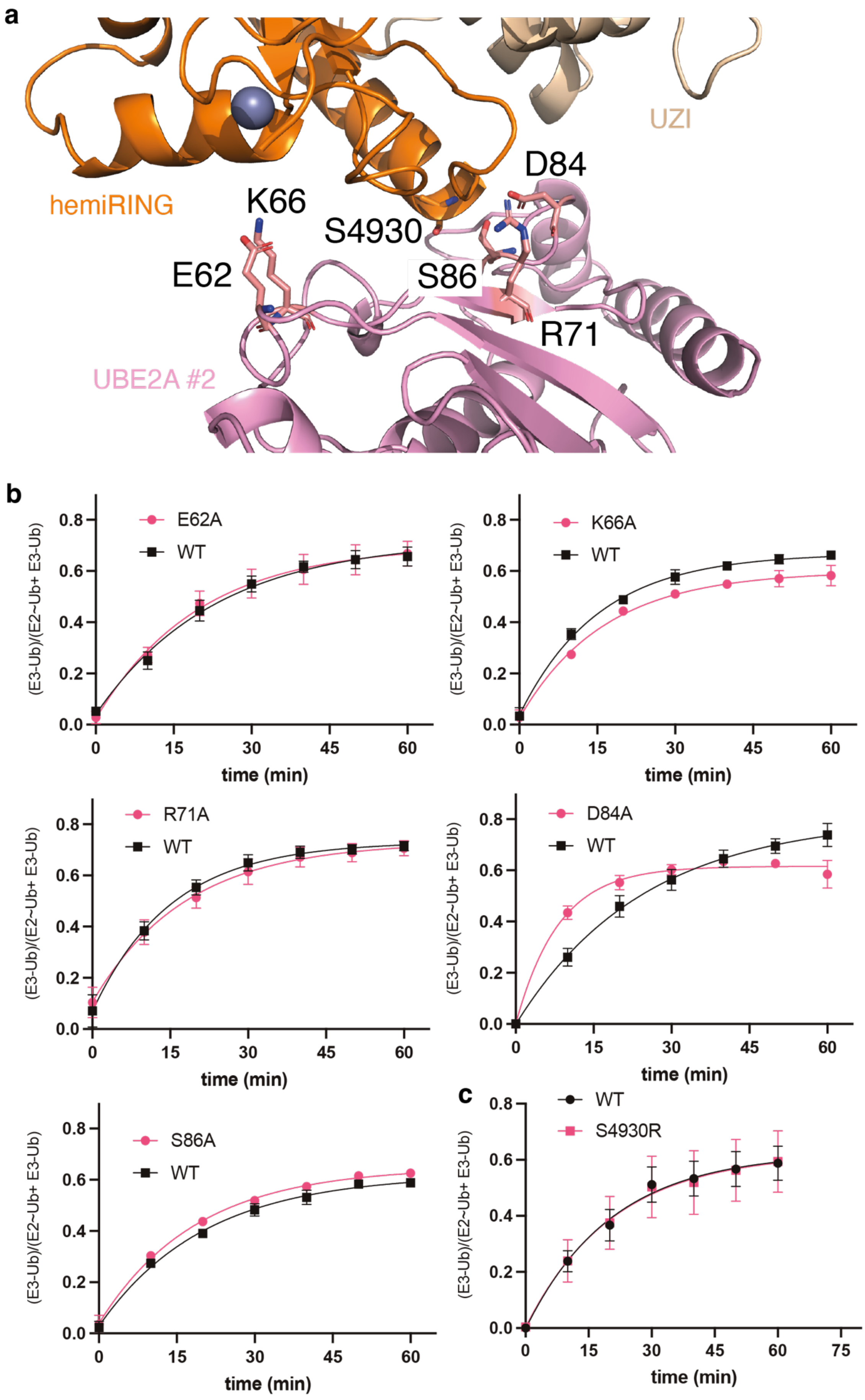
a
hemiRING
UZI
K66
S4930
D84
E62
S86
R71
UBE2A #2
b
E62A
WT
K66A
WT
R71A
WT
D84A
WT
S86A
WT
(E3-Ub)/(E2~Ub+ E3-Ub)
time (min)
c
WT
S4930R
0.0
0.2
0.4
0.6
0.8
0
15
30
45
60
75

**Extended Data Figure 10. investigation of the UBR4-UBE2A #2 interface. a)** The asymmetric unit presented with two UBE2A molecules. UBE2A residues Glu62, Lys66, Arg71, Asp84 and Ser86 were identified as being of potential importance for maintaining the interaction with UBE2A #2 (pink). UBR4 Ser9330 packed closely against UBE2A and it was assumed that mutation to arginine would abolish the UBR4-UBE2A 2 interaction. **b)** The significance of the observed UBR4-UBE2A #2 interaction on UBR4 autoubiquitination was assessed by testing UBE2A alanine mutants of the identified residues. **c)** To exclude the possibility that the UBE2A mutations had a general effect on UBE2A function we tested the steric mutation in UBR4 (Ser4930Arg).

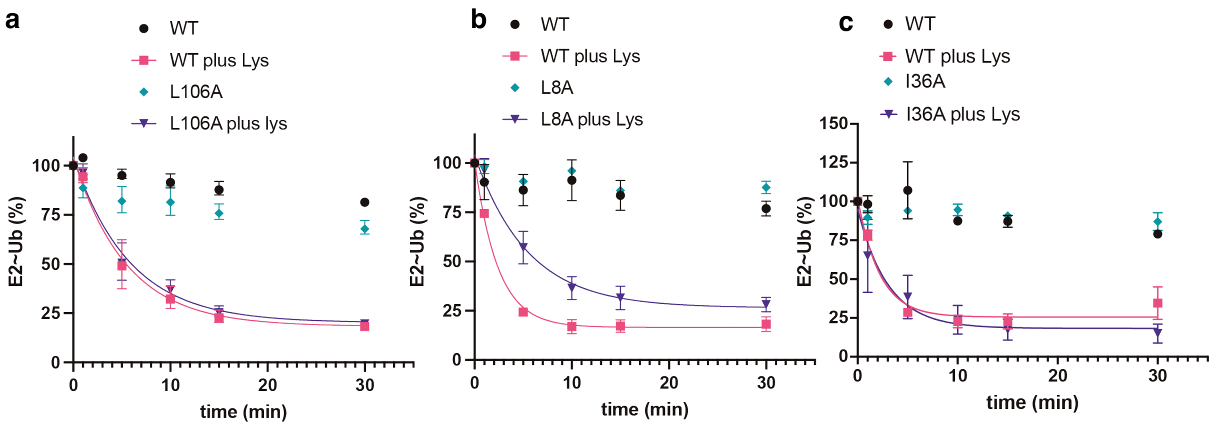

**Extended Data Figure 11. Effect of UBE2A and Ub residues on E3-independent discharge to free lysine deemed important for formation of a closed E2~Ub conformation. a)** UBE2A WT discharges to free lysine (10 mM) with an efficiency comparable to Leu106Ala, indicating this mutation does not impart a general E3-independent defect on UBE2A. **b)** UBE2A Leu8Ala has impaired lysine charge activity indicative of a general defect on E2. **c)** UBE2A Ile36Ala has no discernable effect on E3-independent lysine discharge, consistent with its being distal to the E2 active site and the E2-Ub interface of a closed E2~Ub conformation.

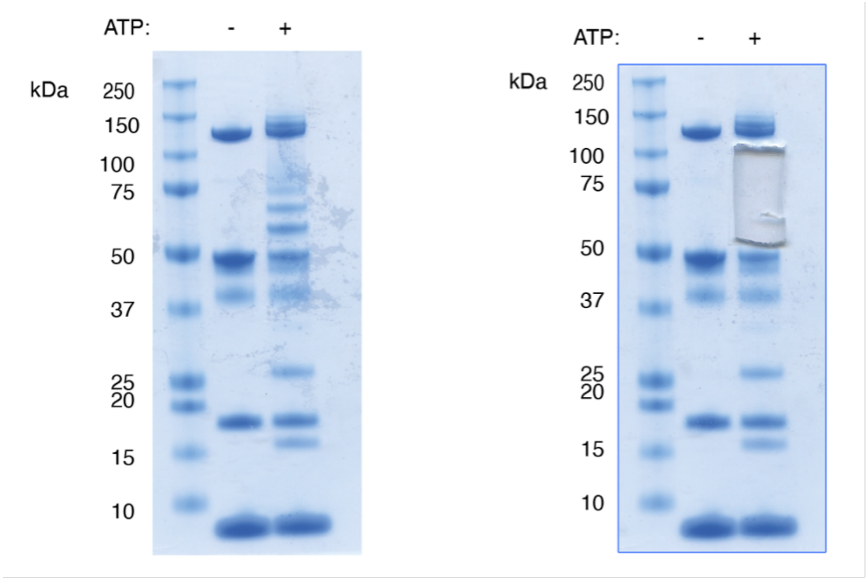

**Extended Data Figure 12. In-gel analysis of autoubiquitination products by mass spectrometry.** An autoubiquitination reaction was carried out with UBR4$_{xtal}$ and the predominant autoubiquitination adducts were excised, dehydrated and resuspended using standard procedures.
